# The Patterns of Alternative TSS Usage Explain the Highly Heterogeneous Landscape of 5’UTR Lengths in Eukaryotes

**DOI:** 10.1101/2024.02.17.580833

**Authors:** Yu Zhan, Zhenbin Hu, Zhaolian Lu, Zhenguo Lin

## Abstract

Transcription start site (TSS) marks the first DNA nucleotide of a gene transcribed into RNA. Accumulating evidence suggests that alternative TSS usage (ATU) is widespread across eukaryotes, in response to environmental changes and tissue-specific needs. However, how ATU is coordinated with changes in gene transcription activity and whether it represents a regulated process or merely transcription initiation errors, remains unclear. To address these questions, we conducted integrative analyses of high-resolution TSS maps and translation efficiency data from multiple eukaryotic organisms. Our results reveal that most ATU events co-occur with differential gene expression and that the direction of ATU is largely consistent with changes in transcription levels. These findings suggest that ATU likely works in concert with transcriptional activity to fine-tune protein production, modulating translation efficiency and mRNA stability through alterations in 5′UTR sequence and structural features. Given its functional importance, the evolution of TSS locations may have been shaped by natural selection, leading to heterogeneous 5′UTR lengths in genes with distinct expression demands. This study offers new insights into the complexity of gene regulation and provides a plausible explanation for the highly variable 5′UTR landscape observed within and among eukaryotic genes.

## Introduction

Gene regulation plays a central role in nearly all cellular processes by controlling the spatial and temporal expression of genes. Dysregulation of gene expression contributes to many human diseases [1]. Most regulatory signals converge at the first step of gene expression, transcription initiation, which determines the location of transcription start sites (TSSs) and transcript abundance [2]. High-throughput sequencing studies have revealed that transcription of most genes is initiated not from a single TSS, but from a cluster of nearby TSSs within core promoter regions [3, 4]. In TSS profiling, these closely spaced sites are grouped into TSS clusters (TCs) to infer core promoter locations [3, 4]. Moreover, genome-wide analyses have shown that most eukaryotic protein-coding genes initiate transcription from multiple, spatially distinct TCs or core promoters [5, 6]. The relative usage of these TCs can shift substantially in response to environmental cues, across tissues, or during development, a phenomenon known as alternative TSS usage (ATU), observed in mammals [7–10], zebrafish [11], budding yeast [5, 12], and pathogenic fungi [13]. ATU has also been implicated in diseases such as cancer [14], neurological disorders, and Alzheimer’s disease [15–18].

ATU can generate transcript isoforms with different translation start codons, producing protein variants with altered or extended N-terminal polypeptides [19–21]. For instance, the human *MAPT* gene, which encodes microtubule-associated protein tau, has a downstream TSS within its second exon [22]. Transcription from this site yields truncated tau proteins with altered localization and function, linked to neurodegeneration [22]. In many other cases, ATU produces transcript isoforms that share the same translation start codon but differ in their 5′ untranslated regions (5′UTRs). Elements and structural features within 5′UTRs can modulate mRNA localization, stability, and translation efficiency (TE) [23–26]. For instance, internal ribosome entry sites (IRES) directly recruit ribosomes to the start codon [27, 28], while while upstream AUGs (uAUGs) can attenuate protein production [29] or generate extended N-terminal isoforms [28]. Genome-wide ribosome density analysis in yeast revealed that the presence of uAUG is significantly associated with poor translation [30]. If a uAUG is followed by a stop codon in the same reading frame, it forms an upstream open reading frame (uORF) that can suppress translation of the downstream main ORF [28, 29, 31–33]. RNA secondary or tertiary structures such as G-quadruplexes can also inhibit ribosomal scanning and initiation [23]. Empirical studies supported that longer 5′ UTRs often reduce TE compared with shorter isoforms [34, 35]. For instance, in breast cancer tissues, BRCA1 transcription frequently initiates from a distal TSS, generating a longer 5′UTR that reduces translation efficiency by tenfold [15]. Stop codons in 5′UTRs may also trigger nonsense-mediated mRNA decay (NMD), shortening transcript half-life [36, 37].

The prevalence of ATU has been interpreted in contrasting ways. Many researchers propose that ATU expands transcriptome diversity, and is maintained by adaptive evolution [3, 38–41]. For instance, analysis of 96 yeast genes showed that 5′UTR isoforms can substantially alter protein output without changing mRNA levels, underscoring the regulatory potential of TSS choice [42]. Others argue that most ATUs result from transcription initiation errors and are non-adaptive [43–45]. A neutral model suggests that variation in 5′UTR length arises primarily through random mutations, with TSS turnover is primarily driven by stochastic processes such as genetic drift [46]. In this model, 5′UTR length is selectively neutral, constrained mainly by the risk of uAUG formation.

Given ATU’s prevalence and potential functional consequences, it is essential to determine whether it represents a regulated mechanism that benefits the organism or merely a byproduct of transcriptional noise [44, 45, 47]. Addressing this question requires quantitative TSS maps that capture both TSS position and transcription abundance. Such maps can be obtained by sequencing the 5′ ends of RNA transcripts and aligning them to a reference genome. Cap Analysis of Gene Expression (CAGE) is one widely used method for this purpose [48, 49]. As CAGE reads mapped to a TSS are theoretically proportional to transcriptional abundance, CAGE-based TSS data can simultaneously quantify 5′ UTR length and transcription abundance for each transcript isoform. Over the past decade, high-resolution TSS maps have been generated for several model organisms using CAGE [5, 50], enabling systematic evaluation of ATU patterns and their functional implications.

In this study, we address this question by systematically interrogating quantitative TSS maps from multiple model organisms, with a focus on *Saccharomyces cerevisiae* [5, 51], as only ∼ 5% of *S. cerevisiae* genes contain introns [52], which minimizes the impacts of other factors such as alternative splicing and using of alternative translation start codon. We present multiple lines of evidence supporting that ATU is likely a regulated process in accordance with gene differential expression, and functions to fine-tune protein output and mRNA stability via 5′UTR variation, supporting a regulated rather than stochastic process.

## MATERIALS AND METHODS

### Generation of quantitative TSS maps based on CAGE data

Raw CAGE sequencing reads from *S. cerevisiae* [5], *S. paradoxus, S. pombe* [51] were downloaded from the NCBI SRA database (Supplementary Table S1). Quantitative TSS maps were generated by aligning raw CAGE reads to the reference genome for each species (Assembly version: sacCer3 for *S. cerevisiae*, N17 for *S. paradoxus*, S. pombe for *S. pombe*) using HISAT2 [53], with the ‘–no-softclip’ option to avoid false-positive TSS calls. The numbers of mapped reads are provided in Supplementary Table S1. Only uniquely mapped reads (MAPQ > 20) were used for TSS identification.

We used the TSSr package [54] to call TSSs from mapped read files. TSS read counts from biological replicates of the same sample were merged for quantification of TSS signals, which were quantified as the numbers of CAGE tags (reads) supporting the TSS per million mapped reads (TPM). Nearby TSSs were grouped as a TSS cluster (TC), representing a core promoter using the ‘peakclu’ function of TSSr (peakDistance = 50, extensionDistance = 25, localThreshold = 0.1, clusterThreshold = 1). Mapped CAGE reads in bam format for human K562 and HepG2 cell lines were downloaded from FANTOM5 [50].

Consensus TCs were generated for the nine *S. cerevisiae* samples, and the two cell lines of human, respectively, by using the ‘consensusCluster’ function in TSSr with an option of ‘dis = 50’. The transcription level of a TC was calculated as the sum of TPM values of all eligible TSSs within it. Only TCs with TPM ≥1 were considered as biologically significant and used for further analyses.

For yeast species, consensus TCs were then assigned to their downstream genes if they are within 750 bp upstream and 50 bp downstream of the start codon of annotated ORFs. TCs associated with non-coding RNA genes in *S. cerevisiae* (e.g., SUTs, XUTs, CUTs, tRNA, rRNA and snoRNAs, Supplementary Dataset S1) were identified using parameters “upstream: 100, upstreamOverlap: 50, downstream: 25”. Consensus TCs in human were assigned to their downstream genes if they were within 1000 bp upstream and 50 bp downstream of the start codon of the gene (ATG).

### Analysis of translation efficiency

To compare translation efficiency (TE) in *S. cerevisiae* under stress (H_2_O_2_) versus control (YPD), ribosome profiling and RNA-seq reads were obtained from Blevins et al. [55]. Ribosome profiling reads were trimmed using fastp [56]. The first and last 10 bases of Read 1, and the first 16 and last 5 bases of Read 2 were removed. Trimmed ribosome profiling reads and RNA-seq reads were aligned to the S. cerevisiae S288c reference genome (R64-2-1) using HISAT2 with max-intron-length = 2500. Reads mapping to rRNA were removed using rRNAdust.

SAM files were converted to BAM format and sorted using samtools [57]. RPF (ribosome-protected fragment) TPM and mRNA TPM were calculated with StringTie [58]. Genes with mRNA TPM < 10 were excluded, and TE was calculated as RPF TPM / mRNA TPM. Average TE values were used for further analyses.

TE and transcript abundance data for mouse fibroblasts (NIH3T3) were obtained from Wang et al. [35]. Transcript abundance and TE values were log₂-transformed for normality. Genes with undefined TE (NA or 0) were excluded.

### Analysis of mRNA half-life data

mRNA decay rate or half-life data for *S. cerevisiae* and *Sch. pombe* were obtained from published datasets [59–62]. In these studies, mRNA was labeled and sequenced at multiple time points following transcriptional inhibition. Decay rates were estimated using a dynamic model [63]. The half-life values was calculated as: half-life = ln(2)/ (decay rate) [64].

### Identification of alternative TSS usage (ATU) and differentially expressed (DE) genes

ATU genes were identified using a paired Wilcoxon test to compare TSS usage between two conditions. To avoid intra-cluster variation, only genes with a dominant TSS distance ≥ 50 bp between conditions were considered. Differentially expressed genes (DEGs) were identified using the deGene function in TSSr [54].

### KEGG pathway and GO enrichment analysis

KEGG pathway gene sets for S. cerevisiae were extracted using the getGenesets function from KEGGREST (org = “sce”, db = “kegg”, cache = TRUE, return.type = “list”), reformatted with the melt function [65], and converted from NCBI Gene ID to KEGG ID. ATU gene enrichment in KEGG pathways was calculated as the observed/expected ratio, with significance assessed by Chi-square test (corrected, *p* < 0.01). Visualization was performed using ggplot2 v3.3.6 [66]. Gene Ontology (GO) enrichment was performed using the Lewis-Sigler Institute GO tools. p-values were computed using the hypergeometric distribution, corrected with Bonferroni adjustment (cutoff p < 0.01) [67].

### Identification of typical structure motifs in 5’UTR

5’UTR sequences were extracted using the “getfasta” tool in bedtools v2.29.2 [68]. Upstream open reading frames (uORFs) were identified using in-house R scripts. Hairpin structures were predicted using ViennaRNA Package [69]. G-quadruplex (G4) motifs were identified as sequences containing ≥ 3 “GGG” tracts [70]. GC content was calculated using in-house scripts.

### Classification of Genes by TSS Cluster Number

Genes were classified as single-cluster (SC) if associated with one TSS cluster, or multi-cluster (MC) if associated with ≥ 2 clusters. Relative variance in expression level or 5′UTR length in *S. cerevisiae* was calculated as SD/mean across nine conditions.

### Calculation of 5′ UTR length and identification of genes with non-optimal 5′ UTR length

The 5′ UTR length of a transcript isoform was calculated as the distance between its dominant TSS, which is the TSS with the largest TPM values within a TC, and the translation start codon of the associated gene. Transcripts with introns in 5′ UTR were excluded from our analysis to avoid potential biases. If a TC’s dominant TSS is located within a gene coding region, which creates a negative 5′ UTR length, the TC was excluded for subsequent analysis. To infer a list of genes with non-optimal 5′ UTR length (outlier gene), a linear regression model for transcript 5′ UTR length and their TPM values was first constructed. Genes falling outside the 95% prediction interval were classified as outliers.

### Evolutionary analysis of *S. cerevisiae* population genomics data

The single nucleotide polymorphism (SNP) dataset was retrieved from whole genome sequencing data of 1,011 *S. cerevisiae* strains [71]. To ensure the integrity of the SNP dataset, multiallelic SNPs were removed using “bcftools” version 1.13 [72]. In our analysis, a promoter SNP was defined as a TSS located within a 750 nucleotide (nt) region upstream of an ORF. The nucleotide diversity (π) for each SNP was computed utilizing “vcftools” version 0.1.16, as outlined in [73]. SNPs with a nucleotide diversity (π) exceeding the threshold of 0.004 were selected for downstream analysis.

### Statistical analysis

All statistical analyses were performed in R v.4.2.1. Statistical analysis information was included in the figure legends. For correlation analysis, the Spearman (ρ) or Pearson (r) correlation coefficients were performed. Binomial tests were employed to determine whether equal numbers of genes in two different patterns. The Wilcoxon test was employed to compare SNP number and π diversity in different gene groups. Scatter plots with density information and boxplots were generated using R packages ggplot2 [66] and ggpointdensity (https://github.com/LKremer/ggpointdensity).

## RESULTS

### Most ATU events coincide with gene differential expression in response to environmental changes

If ATU arises primarily from stochastic transcription initiation errors, its occurrences should be independent of whether and how gene transcription levels change under environmental stimuli. To test this, we identified ATU events in budding yeast *S. cerevisiae* using quantitative TSS maps from nine different growth conditions [5]. TSS maps from cells grown in YPD rich medium were used as a control to identify condition specific ATU events in the remaining eight conditions (Supplementary Dataset S2, Materials and Methods).

To minimize noise from minor transcriptional shifts, we only consider ATU events between two TCs whose dominant TSSs were separated by at least 50 base pairs (bp), as TCs with dominant TSSs < 50 bp apart were merged into a single consensus TC (Supplementary Figure S1A&S2, Table S3, Materials and Methods). In total, we identified 1,157 ATU events from 558 protein-coding genes across the eight conditions. For instance, in YPD medium, transcription of the yeast *MTD1* gene (*YKR080W*) predominantly initiates from its proximal TC (Figure 1A). Upon 2% NaCl treatment, transcription initiation from this proximal TC was nearly abolished, with most transcripts now initiating from the distal TC, representing a clear ATU event in response to environmental stress.

**Figure 1.**
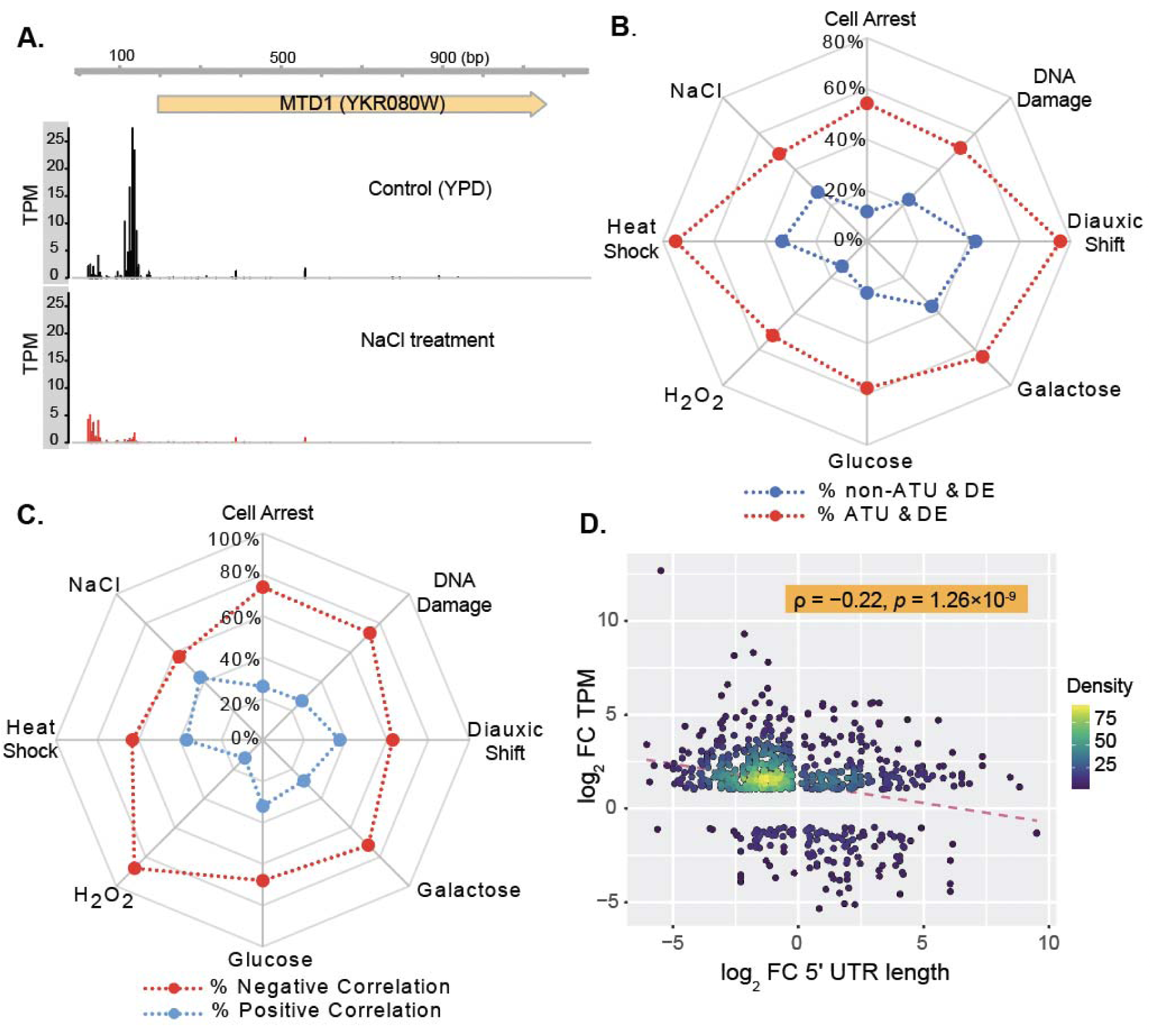
Association between ATU-mediated 5′UTR length changes and gene differential expression under environmental stimuli. **(A)** Example of ATU altering 5′ UTR length by the *MTD1* gene following 1M NaCl treatment via alternative TSS usage (ATU). **(B)** Genes with ATU (red dots) show a higher likelihood of differential expression (DE) compared to genes without ATU (blue dots) across various treatments. Specific treatments are indicated. **(C)** Changes in 5′UTR length by ATU predominantly exhibit a negative correlation with expression level changes; red dots represent genes with negative correlation, blue dots with positive correlation. **(D)** Significant negative correlation between fold changes (FC) in 5′UTR length and gene expression across eight treatments. Transcript density is indicated by color gradient; the red dashed line shows the linear regression fit. Genes with significant transcriptional changes (FC ≥ 2) and ATU events (p < 0.05, Wilcoxon test) between conditions were analyzed.

To infer the extent to which ATU events are associated with gene differential expression (DE), we identified DE genes in each condition from the same dataset. Our data show that 61.71% (714) of all ATU events were associated with significant DE (Supplementary Dataset S2). For example, the ATU in *MTD1* coincided with a marked expression drop from 140.1 TPM in YPD to 33.3 TPM under NaCl treatment. Across all eight treatments, the proportions of genes with ATU that were also DE was significantly higher than for genes without ATU (Figure 1B). In each condition, the number of ATU-associated genes ranged from 109 to 191 (Supplementary Table S4). For example, under heat shock, 141 of 188 genes (75%) with ATU were also DE, a proportion significantly higher than among non-ATU genes.

To determine whether this pattern extends to higher eukaryotes, we analyzed CAGE data from human K562 and HepG2 cell lines [50], because each contains multiple biological replicates that are highly consistent with each other (r ≥ 0.99). Similar results were obtained from both human cell lines, implying that the ATU–DE association is conserved across diverse eukaryotic lineages (Supplementary Figure S1B and Dataset S3). Together, these results demonstrate that ATU events frequently co-occur with differential gene expression, supporting the view that ATU is coordinated with transcriptional regulation rather than arising from random initiation errors.

### The direction of TSS shift aligns with transcriptional changes

In ATU events, transcription initiation shifts either to a more proximal or distal TC, producing transcript isoforms with shortened or elongated 5′UTRs, respectively. We investigated whether the direction of these shifts aligns with changes in gene expression.

Two patterns emerged when ATU co-occurs with differential expression (DE). The first pattern involves transcriptional upregulation associated with a shift to a more proximal promoter, or downregulation associated with a shift from proximal to distal TC. This pattern can be illustrated by the example of the *MTD1* gene (Figure 1A). This pattern represents a negative correlation between transcriptional changes and 5′UTR length. In contrast, the other pattern links transcriptional upregulation with elongated 5′UTRs, reflecting a positive correlation between expression changes and 5′UTR length.

Across all examples examined, 68.07% of ATU–DE coupled events (486 of 714) displayed the *MTD1*-like pattern, indicating a predominant negative correlation between transcriptional change and 5′UTR length. This trend was consistent across individual treatments in *S. cerevisiae* (Figure 1C, Supplementary Dataset S2), and was also observed between two human cell lines (Supplementary Figure S1C, Dataset S3), suggesting that the coordination between TSS shift direction and transcriptional regulation is evolutionarily conserved.

Considering that the distances between TCs of a gene range from 50 bp to several hundred bp (Supplementary Dataset S2), the resulting variation in 5’UTR length between transcript isoforms created by ATU may vary substantially among genes. We then evaluated if there are quantitative impact of changes in 5’UTR length on gene differential expression. We found a significant negative correlation between log₂ fold changes in transcript abundance and 5′UTR length in yeast (ρ = –0.22, *p* = 1.26×10^-9^, Figure 1D), and human cells (ρ = –0.27, *p* = 7.12×10^-16^, Supplementary Figure S1D and Dataset S3). The observation that larger shifts in TSS position, and thus 5′UTR length, correspond to stronger transcriptional changes, supporting a quantitative role for ATU in modulating gene expression via 5′UTR architecture.

### ATU modulates protein production by generating 5’UTR isoforms that influence translation efficiency and mRNA stability

We demonstrate that ATU generating different 5’UTR isoforms frequently coincide with changes in transcription activity, and that 5′UTR length and transcription levels exhibit a negative correlation, suggesting coordinated regulation rather than transcriptional noise. While gene-specific transcription factors modulate protein production via transcriptional control, 5′UTR sequence and structural elements also impact translation efficiency (TE) [24–26, 35]. Thus, ATU-driven variation in 5′UTR isoforms likely fine-tunes protein output by altering TE (Figure 2A), explaining its frequent co-occurrence with transcriptional changes and directional bias.

**Figure 2.**
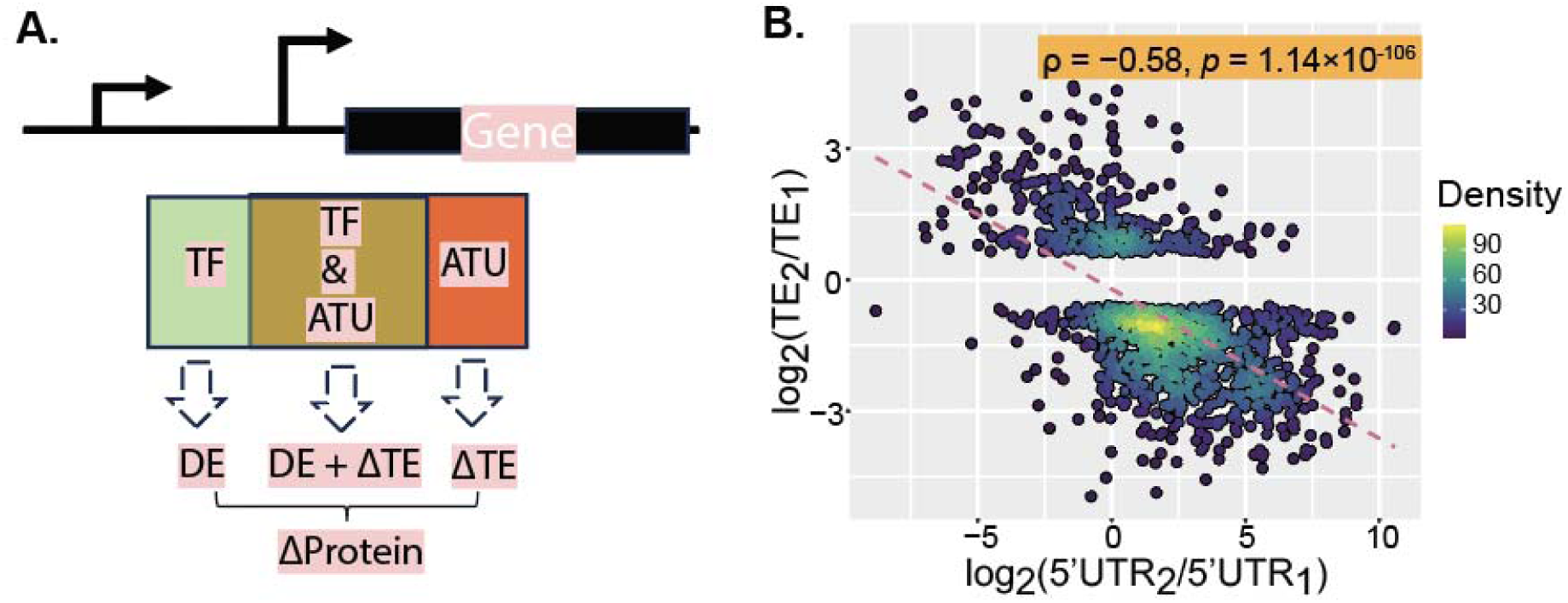
ATU regulates protein production by altering 5′UTR length, impacting translation efficiency. **(A)** Schematic illustration of ATU’s role in modulating gene expression through 5′UTR variation. **(B)** Negative correlation between differences in 5′UTR length and translation efficiency (TE). Differences in 5′UTR length were calculated as log2(5′UTR_₂_ / 5′UTR_₁_), and differences in TE as log2(TE_₂_ / TE_₁_). Only transcript isoform pairs with at least a 1.5-fold difference in TE (|TE_₂_ / TE_₁_| ≥ 1.5) were included.

Supporting this, analysis of *S. cerevisiae* under oxidative stress (H_2_O_2_) revealed that 88.5% of genes with longer 5′UTRs show reduced TE (Supplementary Figure S3A). Similarly, in mouse fibroblasts [35], 79.6% of shorter 5′UTR isoforms associate with higher TE (Supplementary Figure S3B and Dataset S5), with a strong negative correlation between 5′UTR length and TE (ρ = -0.58, *p* = 1.14×10^-106^, Figure 2B and Supplementary Dataset S5).

The influence of 5′UTR length on TE is likely mediated by changes in regulatory elements. In *S. cerevisiae*, 68.25% (4,729 out of 6,929) of 5’UTR differ in upstream open reading frame (uORF) count, which strongly correlates positively with 5′UTR length (ρ = 0.78, *p* < 9.88×10^-323^, Fig. 3A) and negatively with transcript abundance (ρ = –0.56*, p* < 9.88×10^-323^, Fig. 3B). These results suggest that the functional impact of changes in 5’UTR length on TE is likely mediated by changing uORF count. Among isoforms with unchanged uORF counts, secondary structures such as hairpins also vary extensively, with intra-strand base pairs correlating positively with 5′UTR length (ρ = 0.88, *p* < 9.88×10^-323^, Fig. 3C and Dataset S6) and negatively with transcript abundance (ρ = –0.49, *p* < 9.88×10^-323^, Fig. 3D). Similar trends hold for minimum free energy (MTER) and log2 fold changes in transcript abundance (ρ = 0.50, *p* < 9.88×10^-323^, Supplementary Fig. S4A).

**Figure 3.**
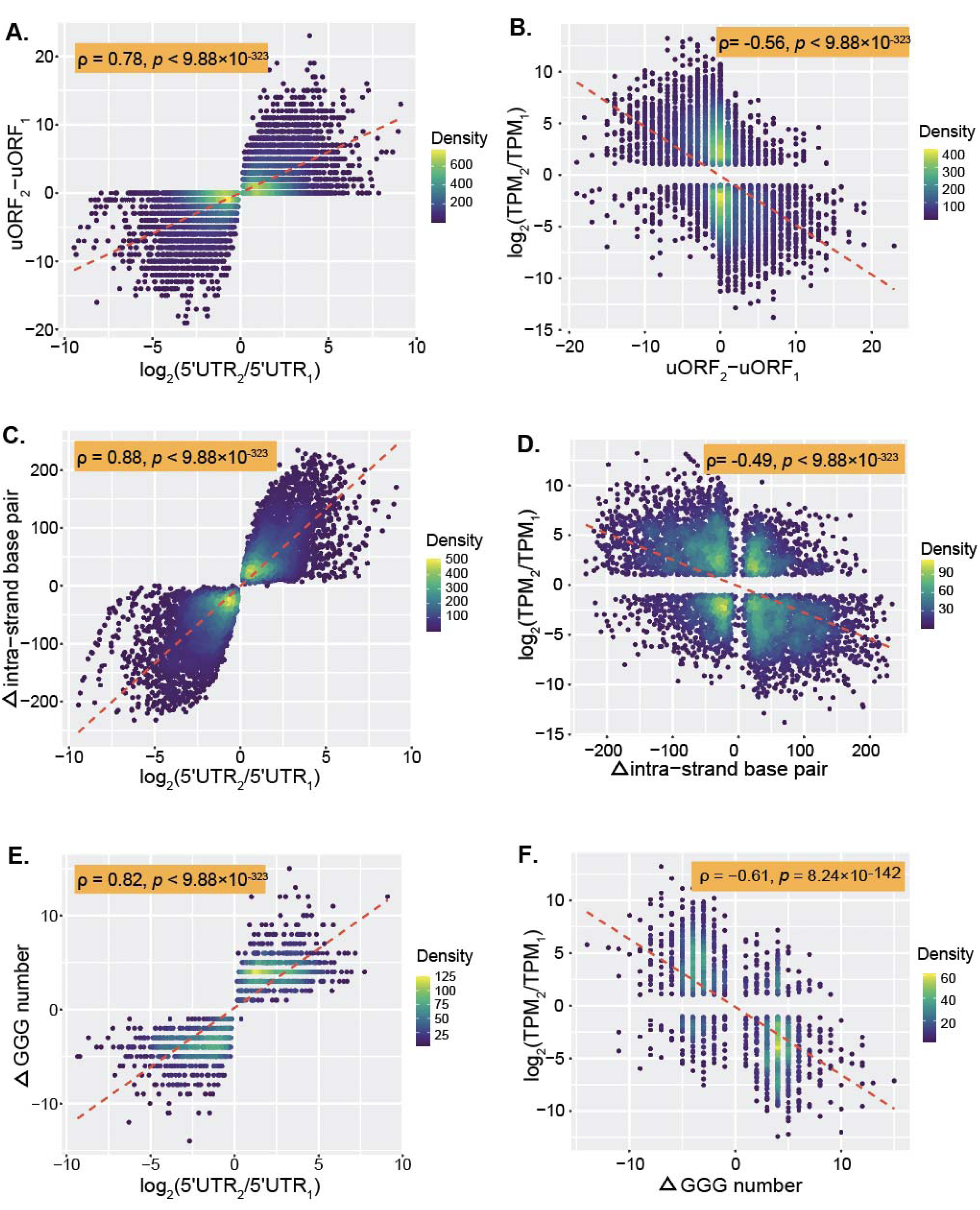
ATU influences functional 5′ UTR motifs: uORFs, hairpins, and G-C-rich sequences. (**A)** Positive correlation between log2 ratio of 5′UTR length [x = log2(5′UTR_₂_ / 5′UTR_₁_)] and change in uORF number [y = uORF_₂_ − uORF_₁_] across transcript isoforms from the same gene. **(B)** Negative correlation between change in uORF number [x = uORF_₂_ − uORF_₁_] and peak expression level [y = log2(TPM_₂_ / TPM_₁_)] from transcript isoforms of the same gene. **(C)** Positive correlation between log2 ratio of 5′UTR length [x = log2(5′UTR_₂_ / 5′UTR_₁_)] and change in intra-strand base pairs [y = base pairs_₂_ − base pairs_₁_] from transcript isoforms of the same gene. **(D)** Negative correlation between change in intra-strand base pairs [x = base pairs_₂_ − base pairs_₁_] and peak expression level [y = log2(TPM_₂_ / TPM_₁_)] from transcript isoforms of the same gene. **(E)** Positive correlation between log2 ratio of 5′UTR length [x = log2(5′UTR_₂_ / 5′UTR_₁_)] and change in GGG motif count [y = GGG_₂_ − GGG_₁_] from transcript isoforms of the same gene. **(F)** Negative correlation between change in GGG motif count [x = GGG_₂_ − GGG_₁_] and fold change in transcript abundance [y = log2(TPM_₂_ / TPM_₁_)] from transcript isoforms of the same gene.

Additional motifs, including G-quadruplexes (G4), which physically block ribosome scanning from the 5’ cap to the start codon [70], occur in ∼10% of yeast 5′UTRs. G4 abundance correlates positively with 5′UTR length (ρ = 0.82, *p* < 9.88×10^-323^, Fig. 3E and Dataset S6) and negatively with transcript levels (ρ = –0.61, *p* = 8.24×10^-142^, Fig. 3F and Dataset S6), indicating another mechanism by which ATU modulates translation. The positional shift of the first GGG motif also negatively correlates with expression (ρ = –0.56, *p* = 1.23×10^-115^, Supplementary Fig. 3B). GC content was reported positively correlated with 5′ UTR length [74]. Consistently, our data show that GC content has a modest positive correlation with 5′ UTR length (ρ = 0.14, *p* = 2.07×10^-26^, Supplementary Fig. S4C and Dataset S6), and a modest negative correlation with transcript abundance (ρ = –0.15, *p* = 1.58×10^-28^, Supplementary Fig. S4D and Dataset S6).

Beyond translation, uORFs harboring premature stop codons can trigger nonsense-mediated decay (NMD) [36, 37], affecting mRNA stability. To test this hypothesis, we retrieved published mRNA half-life datasets for both *S. cerevisiae* and *Schizosaccharomyces pombe* [59–62]. Consistent with this, 5′UTR length negatively correlates with mRNA half-life in both *S. cerevisiae* and *S. pombe* (Supplementary Figs. S5A-C and Dataset S7-S8), indicating that ATU also modulates transcript abundance by influencing mRNA decay.

### A gene’s transcript isoform with shorter 5’UTR tends to have higher transcription level

Our data suggest that shorter 5′UTR isoforms generally have higher translation efficiency and increased mRNA stability, implying that genes favor shorter 5′UTRs to achieve higher expression. We therefore hypothesized that, for most genes, their shorter 5′UTR isoforms should display higher transcription levels than their longer counterparts..

To test this, we analyzed transcript isoform expression and corresponding 5′UTR lengths across multiple yeast growth conditions (Supplementary Dataset S9). In YPD medium, 2,785 TCs from 1,219 genes were identified (Supplementary Dataset S10). Among these, 1,003 shorter isoforms from 736 genes had significantly higher TPM values than their longer isoforms, whereas only 598 shorter isoforms from 445 genes did not follow this pattern (Supplementary Figure S6).

We further examined the quantitative relationship between differences in 5′UTR length and transcription levels. As illustrated by the *UPT5* gene (*YDR398W*), its transcription can be initiated from two TCs that are ∼350 bp apart: the distal TC C1 and the proximal TC C2 (Figure 4A). The difference in transcript levels between C1 and C2 isoforms was quantified as *T* = log_2_(TPM_2_/TPM_1_), where TPM_1_ and TPM_2_ are the transcription level of C1 and C2 isoforms, respectively. The difference in 5’UTR lengths was quantified as *L* = log_2_(5’UTR_2_/5’UTR_1_). Across all genes, we observed a significant negative correlation between T and L (ρ = –0.34, p = 3.77 × 10⁻) (Figure 4B), indicating that shorter 5′UTRs quantitatively associate with higher expression.

**Figure 4.**
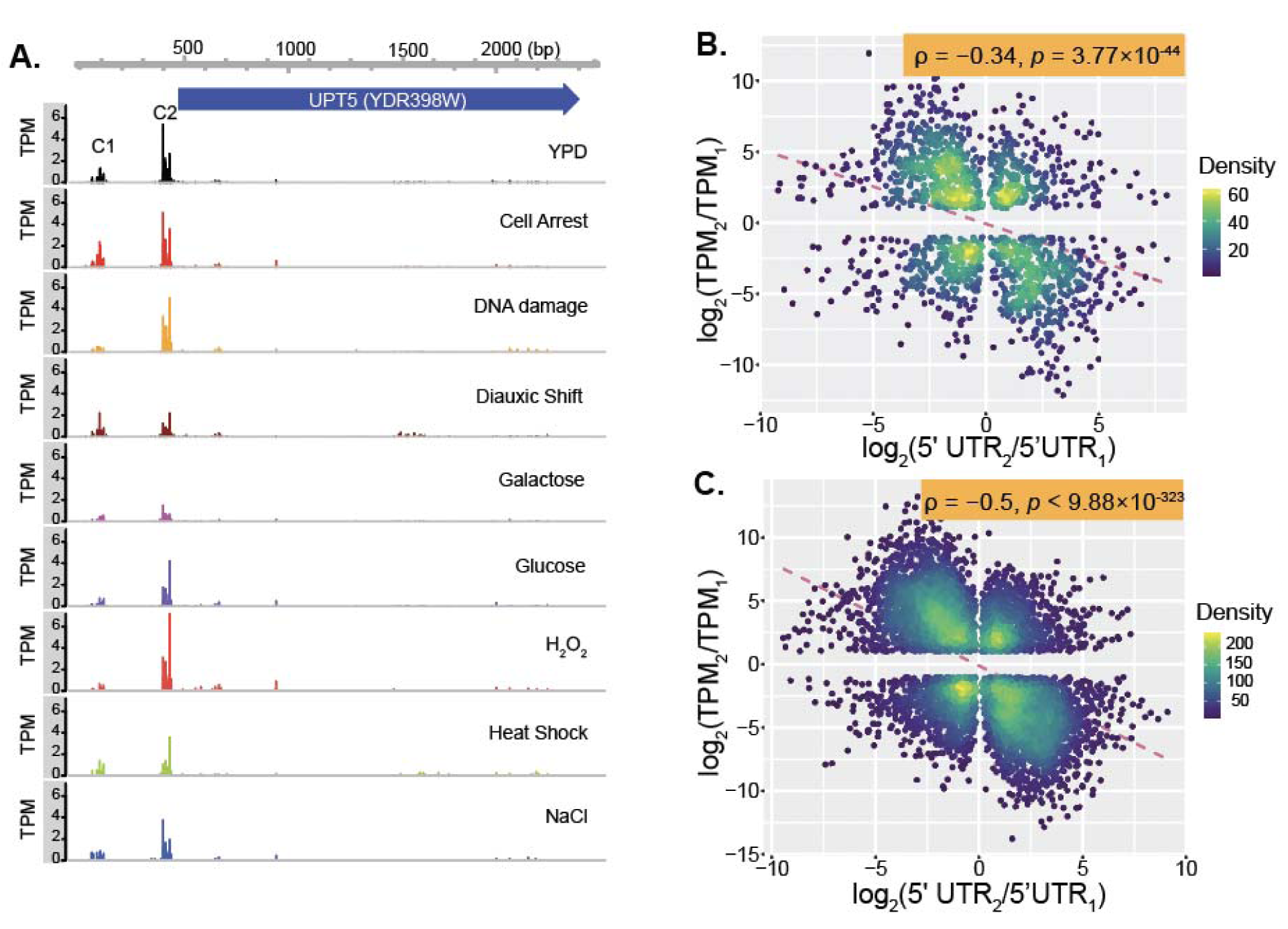
Higher expressed transcript isoforms tend to be equipped with shorter 5’UTR. **(A)** TSS maps of the *UPT5* gene in *S. cerevisiae* across nine growth conditions. The coding sequence (CDS) is shown as a blue arrow. Expression levels of two transcription clusters (TCs) vary significantly across conditions. **(B)** Negative correlation between log2 ratio of 5′UTR lengths [x = log2(5′UTR_₂_ / 5′UTR_₁_)] and relative expression levels [y = log2(TPM_₂_ / TPM_₁_)] for isoforms from the same gene under YP condition. Each dot represents one gene. **(C)** Negative correlation between log2 ratio of 5′UTR lengths [x = log2(5′UTR_₂_ / 5′UTR_₁_)] and peak expression levels [y = log2(TPM_₂_ / TPM_₁_)] across all tested conditions. Peak TPM values represent the highest expression observed for each TC. Each dot represents one gene.

To address potential bias from inactive distal TCs under some conditions, we used peak expression values, the highest TPM observed for each TC across all nine growth conditions, as proxies for maximal activation. For *UPT5*, the distal TC peaked at 10.8 TPM under α-factor arrest, while the proximal TC peaked at 22.3 TPM under H_₂_O_₂_ stress (Figure 4A). Across 9,873 transcript isoforms from all conditions (Supplementary Dataset S11), we performed 6,929 pairwise comparisons among 2,672 genes with multiple isoforms (Supplementary Dataset S12). We found that 67.5% of shorter 5′UTR isoforms had higher peak TPM than longer isoforms (binomial test, *p* < 2.2 × 10^⁻¹^) (Supplementary Fig. S6). This yielded a stronger negative correlation between log2 ratios of 5′UTR length and expression (ρ = –0.5, *p* < 9.88 × 10^⁻³²³^) (Figure 4C). Together, these results reveal an intrinsic link between TSS selection, 5′UTR length, and gene expression demands.

### Genes with shorter 5’UTR tend to have higher expression levels genome-wide

We demonstrate that shorter 5′UTR isoforms generally exhibit higher expression than their longer counterparts. Because gene expression levels vary widely across the genome due to differing protein demands, a negative correlation between 5′UTR length and expression would support the functional importance of 5′UTR length in gene regulation. To mitigate the impact of condition-specific regulation, we calculated peak expression values, the highest TPM observed across nine growth conditions, for 9,873 transcript isoforms in *S. cerevisiae* based on their largest TPM values across nine growth conditions (Supplementary Dataset S11). We found a strong negative correlation between 5′UTR length and peak expression (ρ = −0.5, *p* < 9.88×10^-323^) (Figure 5A), supporting that transcripts with higher expression levels tend to have shorter 5′UTRs both within and across genes.

**Figure 5.**
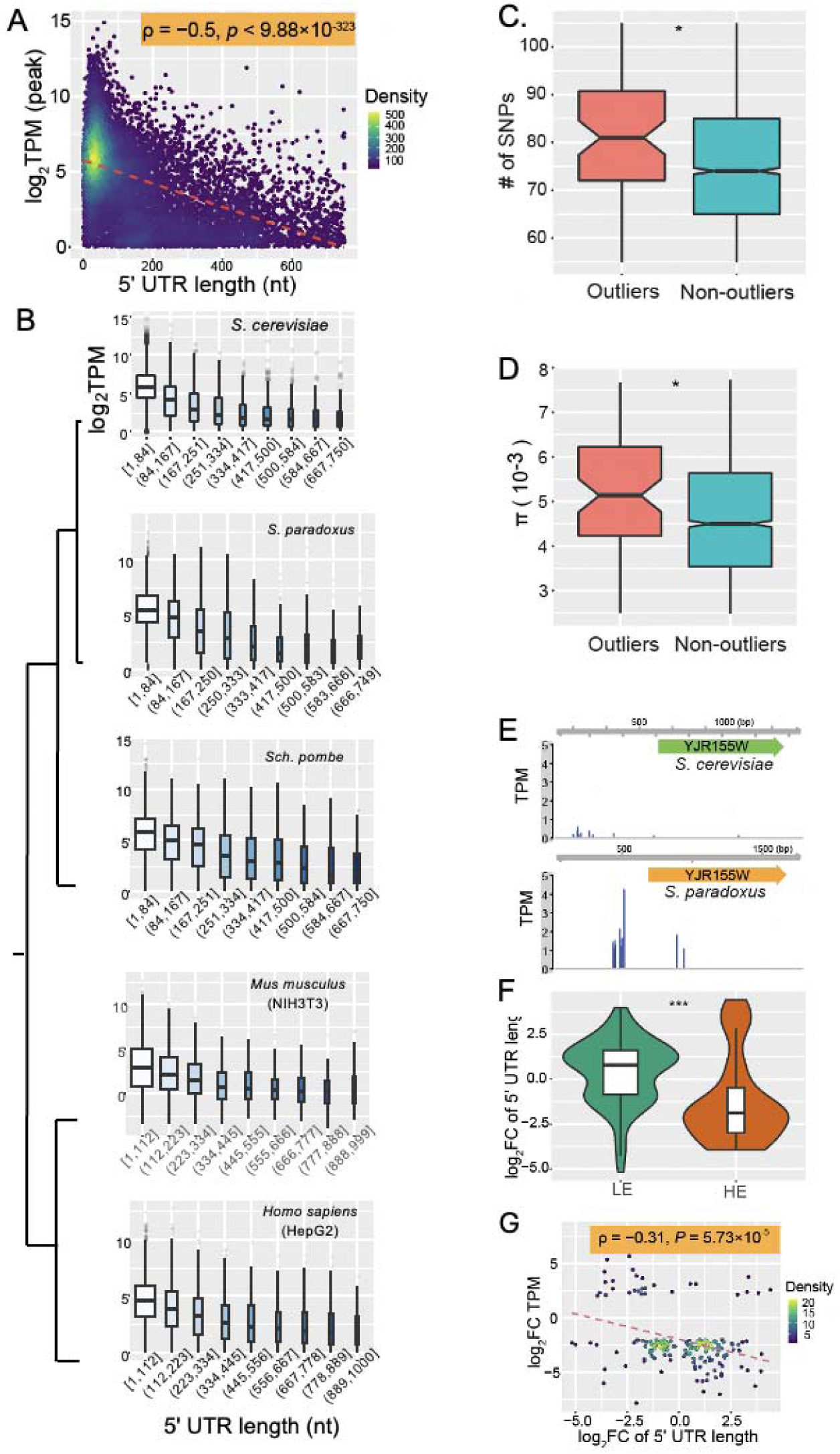
Transcript isoforms with higher expression tend to have shorter 5′UTRs. **(A)** Negative correlation between transcript 5′UTR length and peak expression level in S. cerevisiae. Peak expression is defined as the highest TPM across nine growth conditions. Each dot represents a transcript isoform; color indicates transcript density. The red dashed line shows the linear regression fit. **(B)** Boxplots showing the negative relationship between gene 5′UTR length and expression level across five species. Genes are divided into nine bins based on 5′UTR length. **(C)** Boxplots comparing the number of SNPs in promoter regions of genes with non-optimal 5′UTR lengths (“outliers”) versus other genes. **(D)** Boxplots comparing nucleotide diversity (π) in promoters of outlier genes versus others. *p < 0.05 (Wilcoxon test). **(E)** Example of evolutionary divergence in TSS location and gene expression between *S. cerevisiae AAD10* and its ortholog in *S. paradoxus*, showing differences in 5′UTR length and expression levels. **(F)** Orthologous gene pairs with lower expression (LE) tend to have longer 5′UTRs (log2 FC > 0), while those with higher expression (HE) tend to have shorter 5′UTRs (log2 FC < 0). **(G)** Negative correlation between evolutionary changes in 5′UTR length and gene expression divergence between *S. cerevisiae* and *S. paradoxus*. Color indicates gene density; the red dashed line represents the linear regression fit.

To evaluate the generality of this pattern, we repeated the analysis in four additional species: *S. paradoxus*, *S. pombe*, mouse fibroblasts (NIH3T3), and human HepG2 cells. In all cases, shorter 5′UTRs were consistently associated with higher transcription levels (Figure 5B, Supplementary Dataset S13), confirming that 5′UTR length is a strong, conserved predictor of gene expression across eukaryotes.

### Genes with non-optimal 5′UTRs harbor more promoter SNPs

While 5’UTR generally correlates inversely with gene expression, some genes deviate from this pattern. This may result from a lack of activation signals under tested conditions or from the absence of an optimal TSS within their promoters. Since TSS gain or loss arises from random mutations and optimal 5′UTR length may be constrained by evolutionary chance, according to the theory of mutation-driven evolution [75], we hypothesized that genes with non-optimal 5′UTRs tolerate more promoter mutations than genes with optimal 5′UTRs.

To test this, we used a linear regression model relating 5′UTR length (L) to peak expression (log2TPM): log_2_TPM = −0.0095×*L*+6.46, where *L* is the 5′ UTR length (Supplementary Figure S7 and Dataset S14). Genes whose observed expression fell outside the 95% prediction interval were classified as outliers, and we identified 97 such genes (Supplementary Dataset S14). Functional enrichment analysis revealed that these genes are primarily enriched in signaling pathways (biological processes) of unknown function (Supplementary Fig. S8A), while the remaining genes are predominantly enriched in fundamental biological signaling pathways (Supplementary Fig. S8B).

To assess mutational tolerance, we analyzed promoter SNP data from over 1,000 *S. cerevisiae* isolates [71]. To avoid bias from overlapping promoters, only genes with a single core promoter were included. Outlier genes had a median of 81 SNPs (minor allele frequency ≥ 0.001) in their promoters, significantly higher than the median 74 SNPs for other genes (*p* = 0.01, Wilcoxon test; Figure 5C, Dataset S14). Assuming uniform mutation rates, this elevated SNP density suggests weaker purifying selection on outlier promoters. Supporting this, nucleotide diversity (π) in outlier promoters was also higher (median π = 5.14 × 10⁻³) than in others (π = 4.49 × 10⁻³; *p* = 0.03, Wilcoxon test; Figure 5D). These results indicate that promoters of genes with non-optimal 5′UTR lengths experience relaxed selective constraint, permitting greater accumulation of polymorphisms.

### Evolutionary divergence of 5’UTR length supports its importance in gene expression

Changes in a gene’s 5′UTR length result from gain or loss of TSS within promoter regions. If TSS position, and thus 5′UTR length, is critical for optimal gene expression, evolutionary shifts in TSS locations are unlikely to be random; rather, they should correlate with expression divergence that supports species adaptation to distinct environments.

To test this, we analyzed 4,872 one-to-one orthologous genes between two closely related yeast species, *S. cerevisiae* and *S. paradoxus* (Supplementary Dataset S15). Using CAGE data from cells grown in rich YPD medium [5, 51], we focused on ortholog pairs differing by at least 50 bp in 5′UTR length and exhibiting expression differences of |log2 fold change| ≥ 2.

Consistently, genes with shorter 5′UTRs tend to show higher expression than their orthologs with longer 5′UTRs. For example, *S. cerevisiae AAD10* (*YJR155W*), encoding a putative aryl-alcohol dehydrogenase, is expressed at 2.23 TPM with a 332-nt 5′UTR, whereas its *S. paradoxus* ortholog exhibits higher expression (17.15 TPM) with a markedly shorter 5′UTR (54 nt) (Figure 5E, Supplementary Dataset S15).

At the genome scale, orthologs with lower expression are significantly more likely to have longer 5′UTRs compared to their higher-expressed counterparts (Figure 5F, *p* < 0.001, t-test). Moreover, the extent of 5′UTR length divergence negatively correlates with gene expression divergence (ρ = −0.31, *p* = 5.73 × 10⁻) (Figure 5G).

These observations are consistent with isoform-level patterns within species, and further support the notion that proper TSS location, and resulting 5’UTR length, is a key evolutionary mechanism for fine-tuning gene expression.

### Natural selection shapes TSS locations to meet gene expression demands

Our findings support the importance of TSS positioning and 5’UTR length in regulating gene expression. Based on this, we propose a model in which natural selection shapes TSS locations to optimize expression according to gene-specific demands (Fig. 6A). Because TSSs can arise or be lost via single mutations in promoter regions (Figs. 6B-D) [51], stochastic mutations generate transcript isoforms with varying 5′UTR lengths. The evolutionary fate of a novel TSS depends largely on its position relative to the gene’s expression requirements.

**Figure 6.**
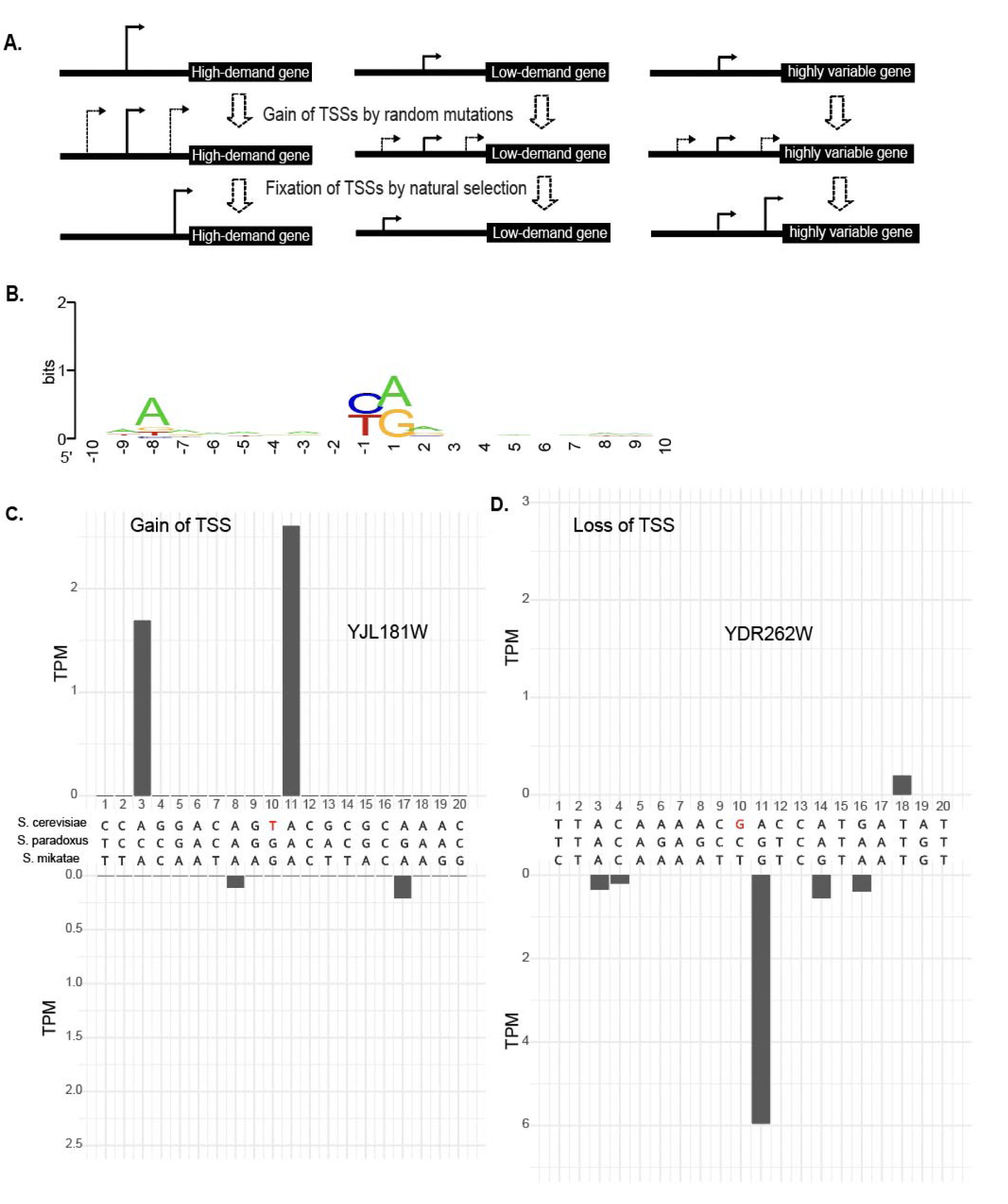
An evolutionary model of TSS explains the heterogeneity of 5′UTR length within and between genes. (A) Genes may gain or lose TSSs through stochastic mutations. The evolutionary fate of a new TSS is largely determined by its genomic location and the gene’s expression demand. For high-demand genes, TSSs generating shorter 5′UTRs, which confer higher translation efficiency, are favored by natural selection and become fixed. Conversely, for genes with lower expression demand, TSSs producing longer 5′UTRs are favored to reduce translation efficiency and protein output. **(B)** Typical sequence features near a TSS (−10 to +10 bp) include a pyrimidine at position -1 and a purine at position +1. **(C)** Example of TSS gain through a G-to-T mutation at position +10, leading to the emergence of a TSS at position +11 in the *YJL181W* gene of *S. cerevisiae*. The upper panel shows TSS expression in *S. cerevisiae*, the lower panel in *S. mikatae*, and the middle section depicts a sequence alignment of the region across three species. **(D)** Example of TSS loss through a C-to-G mutation at position +10, resulting in the loss of a TSS at position +11 in the *YDR262W* gene of *S. cerevisiae*.

For genes with high expression demand, such as ribosomal protein genes, selection favors proximal TSSs that produce shorter 5′UTRs, enhancing translation efficiency and mRNA stability. Such alleles confer a fitness advantage and tend to be fixed in populations. Conversely, genes with low expression demand favor distal TSSs that generate longer 5′UTRs, reducing translation and facilitating repression. For highly variable genes that require dynamic expression across conditions, acquiring multiple TSSs increases transcript diversity, enabling flexible regulation via alternative transcriptional usage (ATU).

Our model predicts that genes with higher expression demand generally possess shorter 5′UTRs. This is supported by the observed negative correlation between expression levels and 5′UTR length (Fig. 5A). To further test this prediction, we examined the distributions of 5′ UTR lengths among genes of different functional categories in *S. cerevisiae*. Gene functional classifications were obtained based on annotation information from Gene Ontology (GO) [76] and Kyoto Encyclopedia of Genes and Genomes (KEGG) [77]. Our results show that 5′UTR lengths vary significantly among functional categories in *S. cerevisiae* (Supplementary Fig. S9A-B, ANOVA, *p* < 0.001). For example, ribosomal protein genes have a median 5′UTR length of 28 nt, markedly shorter than the 128 nt median of “MAPK signaling pathway” genes, which respond to diverse stimuli. These findings are consistent with prior reports linking 5′UTR length to gene function [78].

Our model also predicts that genes with greater expression variability across different growth conditions are more likely to harbor multiple core promoters to increase transcript and expression diversity. Transcription factor (TF) genes, known for dynamic regulation, can be used to test this prediction. Analyzing TF genes from YEASTRACT [79], we found 65.8% contain multiple TCs, significantly higher than 48.1% in non-TF genes and 51.6% in housekeeping ribosomal genes (Supplementary Dataset S16). On average, TF genes produce 2.3 distinct 5′UTR isoforms, compared to 1.8 and 1.5 in non-TF and ribosomal genes, respectively.

Finally, we compared genes with multiple core promoters (MC genes; n = 2,672) to those with a single core promoter (SC genes; n = 2,839). MC genes exhibit significantly greater expression variability and 5′UTR length variance than SC genes (Supplementary Figure S10, Dataset S16), highlighting that multiple promoters confer regulatory flexibility in response to environmental and cellular changes.

## DISCUSSIONS

### ATU as an additional layer of gene regulation

Our study demonstrate that ATU in eukaryotes is a non-random, regulated process closely linked to gene expression changes in response to environmental stimuli. Rather than being transcriptional noise, ATU modulates gene expression outcomes by altering 5′UTR length, thereby influencing translation efficiency (TE) and mRNA stability.

This supports prior speculation that ATU adds an extra regulatory layer to fine-tune gene expression, contributing to the complexity of gene regulation [35, 80].

While TFs often serve as binary switches for gene activation or repression [1], ATU provides more nuanced control. For example, using a distal TSS can reduce protein output by lowering TE without altering transcription levels. Importantly, ATU and TF-mediated regulation are complementary: TFs may differentially activate promoters to induce alternative TSS usage, as seen in genes like *Runx1*[81].

We also show that changes in protein production can occur without changes in transcription level (Figure 2A). About 60% of ATU events coincide with differential expression, but many do not. Even without transcriptional changes, shorter 5′UTR isoforms often exhibit higher TE, as observed in mouse fibroblasts where ∼80% of shorter 5′UTR isoforms translate more efficiently (Supplementary Figure S3B). This reinforces ATU’s role as a regulatory layer complementary to TF-driven transcriptional control.

Though most genes display a negative correlation between 5′UTR length and expression, some exhibit the opposite pattern (Fig. 1D & 5A, Supplementary Figure 5). Short 5′UTRs can cause ribosome overcrowding and collisions, triggering RNA degradation pathways that suppress expression [82, 83], highlighting the mechanistic complexity of gene regulation.

### Functional importance of 5’UTR length

We provide several lines of evidence supporting the importance of 5′ UTR length in gene expression regulation. We find strong negative correlations between 5′UTR length and transcription levels both within and across genes, and evolutionary divergence in 5′UTR length associates with expression divergence. Rao et al. similarly reported a weaker negative correlation (r = −0.14) using chicken EST data [84]. The stronger correlation observed here (ρ = −0.50, Figure 5A) likely reflects our use of high-resolution TSS maps and peak expression values across nine conditions, reducing confounding by condition-specific TF availability. However, since these nine conditions capture only a fraction of possible regulatory contexts, expanded sampling may reveal an even stronger negative correlation.

### A selection-based model explains the diverse landscape of 5′UTR lengths

Although a single nucleotide (nt) of 5’UTR was shown to be technically sufficient for the initiation of translation in mammals based on an *in vitro* study [85], many eukaryotic genes possess unexpectedly long 5′ UTRs, up to several thousand nucleotides in invertebrates [86] and in *S. pombe*, which has a highly compact genome [87]. While long 5′UTRs risk harboring inhibitory elements, their conservation suggests functional importance. A prior null model proposed that stochastic gain and loss of TSSs produce the broad 5′UTR length distribution [88]. Our selection-based model differs by emphasizing non-random retention of distal TSSs shaped by expression demand. Housekeeping genes with high protein demand [89] favor proximal TSSs and short 5′UTRs, while low-demand or stress-responsive genes benefit from distal TSSs generating longer 5′UTRs that repress expression. This adaptive tuning explains the 1,000-fold variation in 5′UTR lengths within genomes.

Our model also well explains the negative correlation between the 5′ UTR length and transcription levels observed in all species examined. Because genes of different functional groups have very different expression demands, our model explains why the 5′ UTR length varies substantially among functional groups of genes [89]. It further explains why many genes lack perfectly optimized 5′UTR lengths due to limited mutational opportunities or compensatory mechanisms [75].

Finally, our model suggests natural selection as the primary force shaping 5′UTR length distributions, rather than genetic drift. The functional impact of TSS-creating variants governs their evolutionary fate, explaining observations such as longer 5′UTRs in species with large populations [90]. It also reconciles discrepancies between null model predictions—like the GC content–5′UTR length relationship—and empirical data [74], by recognizing gene-specific expression demand as the dominant constraint over sequence features like uAUGs.

## Acknowledgments

This work was supported by the US National Science Foundation grant (1951332) and the Saint Louis University 2022 President’s Research Fund to ZG. L.

## Reference

1. Lee, T.I. and R.A. Young, Transcriptional Regulation and its Misregulation in Disease. Cell, 2013. 152(6): p. 1237–1251.

2. Lenhard, B., A. Sandelin, and P. Carninci, Metazoan promoters: emerging characteristics and insights into transcriptional regulation. Nature Reviews. Genetics, 2012. 13(4): p. 233–245.

3. Carninci, P., et al., Genome-wide analysis of mammalian promoter architecture and evolution. Nature Genetics, 2006. 38(6): p. 626–635.

4. Sandelin, A., et al., Mammalian RNA polymerase II core promoters: insights from genome-wide studies. Nat Rev Genet, 2007. 8(6): p. 424–36.

5. Lu, Z. and Z. Lin, Pervasive and dynamic transcription initiation in Saccharomyces cerevisiae. Genome Research, 2019. 29(7): p. 1198–1210.

6. Yamashita, R., et al., Genome-wide characterization of transcriptional start sites in humans by integrative transcriptome analysis. Genome Research, 2011. 21(5): p. 775–789.

7. Pozner, A., et al., Developmentally regulated promoter-switch transcriptionally controls Runx1 function during embryonic hematopoiesis. BMC Dev Biol, 2007. 7: p. 84.

8. Shephard, E.A., et al., Alternative promoters and repetitive DNA elements define the species-dependent tissue-specific expression of the FMO1 genes of human and mouse. The Biochemical Journal, 2007. 406(3): p. 491–499.

9. Steinthorsdottir, V., et al., Multiple novel transcription initiation sites for NRG1. Gene, 2004. 342(1): p. 97–105.

10. Zhang, P., et al., Relatively frequent switching of transcription start sites during cerebellar development. BMC Genomics, 2017. 18: p. 461.

11. Haberle, V., et al., Two independent transcription initiation codes overlap on vertebrate core promoters. Nature, 2014. 507(7492): p. 381–5.

12. Waern, K. and M. Snyder, Extensive transcript diversity and novel upstream open reading frame regulation in yeast. G3 (Bethesda), 2013. 3(2): p. 343–52.

13. Dang, T.T.V., J. Colin, and G. Janbon, Alternative Transcription Start Site Usage and Functional Implications in Pathogenic Fungi. J Fungi (Basel), 2022. 8(10).

14. Demircioglu, D., et al., A Pan-cancer Transcriptome Analysis Reveals Pervasive Regulation through Alternative Promoters. Cell, 2019. 178(6): p. 1465–1477 e17.

15. Sobczak, K. and W.J. Krzyzosiak, Structural determinants of BRCA1 translational regulation. The Journal of Biological Chemistry, 2002. 277(19): p. 17349–17358.

16. Anvar, S.Y., et al., Full-length mRNA sequencing uncovers a widespread coupling between transcription initiation and mRNA processing. Genome Biology, 2018. 19(1): p. 46.

17. Tan, W., et al., Molecular cloning of a brain-specific, developmentally regulated neuregulin 1 (NRG1) isoform and identification of a functional promoter variant associated with schizophrenia. The Journal of Biological Chemistry, 2007. 282(33): p. 24343–24351.

18. Mihailovich, M., et al., Complex translational regulation of BACE1 involves upstream AUGs and stimulatory elements within the 5’ untranslated region. Nucleic Acids Research, 2007. 35(9): p. 2975–2985.

19. Karlsson, K., P. Lönnerberg, and S. Linnarsson, Alternative TSSs are co-regulated in single cells in the mouse brain. Molecular Systems Biology, 2017. 13(5): p. 930.

20. Gacita, A.M., et al., Altered Enhancer and Promoter Usage Leads to Differential Gene Expression in the Normal and Failed Human Heart. Circulation: Heart Failure, 2020. 13(10): p. e006926.

21. Dahale, S., et al., Cap analysis of gene expression reveals alternative promoter usage in a rat model of hypertension. Life Science Alliance, 2022. 5(4): p. e202101234.

22. Huin, V., et al., Alternative promoter usage generates novel shorter MAPT mRNA transcripts in Alzheimer’s disease and progressive supranuclear palsy brains. Sci Rep, 2017. 7(1): p. 12589.

23. Leppek, K., R. Das, and M. Barna, Functional 5′ UTR mRNA structures in eukaryotic translation regulation and how to find them. Nature reviews. Molecular cell biology, 2018. 19(3): p. 158–174.

24. Ding, Y., P. Shah, and J.B. Plotkin, Weak 5′-mRNA Secondary Structures in Short Eukaryotic Genes. Genome Biology and Evolution, 2012. 4(10): p. 1046–1053.

25. Guo, N., et al., Alternative transcription start site selection in Mr-OPY2 controls lifestyle transitions in the fungus Metarhizium robertsii. Nature Communications, 2017. 8(1): p. 1565.

26. Uppala, J.K., et al., The cap-proximal RNA secondary structure inhibits preinitiation complex formation on HAC1 mRNA. Journal of Biological Chemistry, 2022. 298(3): p. 101648.

27. Kozak, M., A second look at cellular mRNA sequences said to function as internal ribosome entry sites. Nucleic Acids Res, 2005. 33(20): p. 6593–602.

28. Hinnebusch, A.G., I.P. Ivanov, and N. Sonenberg, Translational control by 5’-untranslated regions of eukaryotic mRNAs. Science, 2016. 352(6292): p. 1413–6.

29. Dvir, S., et al., Deciphering the rules by which 5’-UTR sequences affect protein expression in yeast. Proc Natl Acad Sci U S A, 2013. 110(30): p. E2792–801.

30. Arribere, J.A. and W.V. Gilbert, Roles for transcript leaders in translation and mRNA decay revealed by transcript leader sequencing. Genome Res, 2013. 23(6): p. 977–87.

31. Lin, Y., et al., Impacts of uORF codon identity and position on translation regulation. Nucleic Acids Res, 2019. 47(17): p. 9358–9367.

32. May, G.E., et al., Unraveling the influences of sequence and position on yeast uORF activity using massively parallel reporter systems and machine learning. Elife, 2023. 12.

33. May, G.E. and C.J. McManus, High-Throughput Quantitation of Yeast uORF Regulatory Impacts Using FACS-uORF. Methods Mol Biol, 2022. 2404: p. 331–351.

34. Araujo, P.R., et al., Before It Gets Started: Regulating Translation at the 5’ UTR. Comp Funct Genomics, 2012. 2012: p. 475731.

35. Wang, X., et al., Pervasive isoform-specific translational regulation via alternative transcription start sites in mammals. Molecular Systems Biology, 2016. 12(7): p. 875.

36. He, F. and A. Jacobson, Nonsense-Mediated mRNA Decay: Degradation of Defective Transcripts Is Only Part of the Story. Annu Rev Genet, 2015. 49: p. 339–66.

37. Kervestin, S. and A. Jacobson, NMD: a multifaceted response to premature translational termination. Nat Rev Mol Cell Biol, 2012. 13(11): p. 700–12.

38. Landry, J.R., D.L. Mager, and B.T. Wilhelm, Complex controls: the role of alternative promoters in mammalian genomes. Trends Genet, 2003. 19(11): p. 640–8.

39. Davuluri, R.V., et al., The functional consequences of alternative promoter use in mammalian genomes. Trends in Genetics, 2008. 24(4): p. 167–177.

40. de Klerk, E. and P.A. t Hoen, Alternative mRNA transcription, processing, and translation: insights from RNA sequencing. Trends Genet, 2015. 31(3): p. 128–39.

41. Policastro, R.A. and G.E. Zentner, Global approaches for profiling transcription initiation. Cell Reports Methods, 2021. 1(5): p. 100081.

42. Rojas-Duran, M.F. and W.V. Gilbert, Alternative transcription start site selection leads to large differences in translation activity in yeast. RNA, 2012. 18(12): p. 2299–305.

43. Xu, C., J.-K. Park, and J. Zhang, Evidence that alternative transcriptional initiation is largely nonadaptive. PLoS biology, 2019. 17(3): p. e3000197.

44. Jinrui Xu, H.E.P., Jill E. Moore, Mark B. Gerstein, and Zhiping Weng, Building integrative functional maps of gene regulation. Human Molecular Genetics, 2022. 31.

45. Forutan, M., et al., Evolution of tissue and developmental specificity of transcription start sites in Bos taurus indicus. Commun Biol, 2021. 4(1): p. 829.

46. Lynch, M., D.G. Scofield, and X. Hong, The evolution of transcription-initiation sites. Molecular Biology and Evolution, 2005. 22(4): p. 1137–1146.

47. Chia, M., et al., High-resolution analysis of cell-state transitions in yeast suggests widespread transcriptional tuning by alternative starts. Genome Biology, 2021. 22(1): p. 34.

48. Murata, M., et al., Detecting expressed genes using CAGE. Methods Mol Biol, 2014. 1164: p. 67–85.

49. Morioka, M.S., et al., Cap Analysis of Gene Expression (CAGE): A Quantitative and Genome-Wide Assay of Transcription Start Sites. Methods Mol Biol, 2020. 2120: p. 277–301.

50. Lizio, M., et al., Gateways to the FANTOM5 promoter level mammalian expression atlas. Genome Biology, 2015. 16: p. 22.

51. Lu, Z. and Z. Lin, The origin and evolution of a distinct mechanism of transcription initiation in yeasts. Genome Research, 2020: p. gr.264325.120.

52. Parenteau, J., et al., Deletion of Many Yeast Introns Reveals a Minority of Genes that Require Splicing for Function. Molecular Biology of the Cell, 2008. 19(5): p. 1932–1941.

53. Kim, D., et al., Graph-based genome alignment and genotyping with HISAT2 and HISAT-genotype. Nat Biotechnol, 2019. 37(8): p. 907–915.

54. Lu, Z., et al., TSSr: an R package for comprehensive analyses of TSS sequencing data. NAR Genomics and Bioinformatics, 2021. 3(4): p. lqab108.

55. Blevins, W.R., et al., Uncovering de novo gene birth in yeast using deep transcriptomics. Nature Communications, 2021. 12(1): p. 604.

56. Chen, S., et al., fastp: an ultra-fast all-in-one FASTQ preprocessor. Bioinformatics, 2018. 34(17): p. i884–i890.

57. Li, H., et al., The Sequence Alignment/Map format and SAMtools. Bioinformatics, 2009. 25(16): p. 2078–9.

58. Pertea, M., et al., StringTie enables improved reconstruction of a transcriptome from RNA-seq reads. Nature Biotechnology, 2015. 33(3): p. 290–295.

59. Harigaya, Y. and R. Parker, The link between adjacent codon pairs and mRNA stability. BMC Genomics, 2017. 18(1): p. 364.

60. Chan, L.Y., et al., Non-invasive measurement of mRNA decay reveals translation initiation as the major determinant of mRNA stability. Elife, 2018. 7.

61. Neymotin, B., R. Athanasiadou, and D. Gresham, Determination of in vivo RNA kinetics using RATE-seq. RNA, 2014. 20(10): p. 1645–52.

62. Hasan, A., et al., Systematic analysis of the role of RNA-binding proteins in the regulation of RNA stability. PLoS Genet, 2014. 10(11): p. e1004684.

63. Miller, C., et al., Dynamic transcriptome analysis measures rates of mRNA synthesis and decay in yeast. Mol Syst Biol, 2011. 7: p. 458.

64. Chen, C.Y., N. Ezzeddine, and A.B. Shyu, Messenger RNA half-life measurements in mammalian cells. Methods Enzymol, 2008. 448: p. 335–57.

65. Wickham, H., Reshaping Data with the reshape Package. Journal of Statistical Software, 2007. 21: p. 1–20.

66. Wickham, H., ggplot2 Elegant Graphics for Data Analysis. 2009.

67. Boyle, E.I., et al., GO::TermFinder—open source software for accessing Gene Ontology information and finding significantly enriched Gene Ontology terms associated with a list of genes. Bioinformatics (Oxford, England), 2004. 20(18): p. 3710–3715.

68. Quinlan, A.R. and I.M. Hall, BEDTools: a flexible suite of utilities for comparing genomic features. Bioinformatics, 2010. 26(6): p. 841–842.

69. Lorenz, R., et al., ViennaRNA Package 2.0. Algorithms for Molecular Biology, 2011. 6(1): p. 26.

70. Capra, J.A., et al., G-quadruplex DNA sequences are evolutionarily conserved and associated with distinct genomic features in Saccharomyces cerevisiae. PLoS Comput Biol, 2010. 6(7): p. e1000861.

71. Peter, J., et al., Genome evolution across 1,011 Saccharomyces cerevisiae isolates. Nature, 2018. 556(7701): p. 339–344.

72. Danecek, P., et al., Twelve years of SAMtools and BCFtools. Gigascience, 2021. 10(2).

73. Danecek, P., et al., The variant call format and VCFtools. Bioinformatics, 2011. 27(15): p. 2156–2158.

74. Reuter, M., et al., A Test of the Null Model for 5′ UTR Evolution Based on GC Content. Molecular Biology and Evolution, 2008. 25(5): p. 801–804.

75. Nei, M., Mutation-driven evolution. 2014, Oxford: Oxford University Press. xi, 244 p., 4 p. of plates.

76. Ashburner, M., et al., Gene ontology: tool for the unification of biology. The Gene Ontology Consortium. Nat Genet, 2000. 25(1): p. 25–9.

77. Kanehisa, M., et al., The KEGG databases at GenomeNet. Nucleic Acids Res, 2002. 30(1): p. 42–6.

78. Spealman, P., et al., Conserved non-AUG uORFs revealed by a novel regression analysis of ribosome profiling data. Genome Res, 2018. 28(2): p. 214–222.

79. Monteiro, P.T., et al., YEASTRACT+: a portal for cross-species comparative genomics of transcription regulation in yeasts. Nucleic Acids Res, 2020. 48(D1): p. D642–D649.

80. Rojas-Duran, M.F. and W.V. Gilbert, Alternative transcription start site selection leads to large differences in translation activity in yeast. RNA, 2012. 18(12): p. 2299–2305.

81. Thomas, A.L., et al., Transcriptional Regulation of RUNX1: An Informatics Analysis. Genes (Basel), 2021. 12(8).

82. Simms, C.L., L.L. Yan, and H.S. Zaher, Ribosome Collision Is Critical for Quality Control during No-Go Decay. Mol Cell, 2017. 68(2): p. 361–373 e5.

83. D’Orazio, K.N. and R. Green, Ribosome states signal RNA quality control. Mol Cell, 2021. 81(7): p. 1372–1383.

84. Rao, Y.S., et al., Relationship between 5′ UTR length and gene expression pattern in chicken. Genetica, 2013. 141(7): p. 311–318.

85. Hughes, M.J. and D.W. Andrews, A single nucleotide is a sufficient 5’ untranslated region for translation in an eukaryotic in vitro system. FEBS Lett, 1997. 414(1): p. 19–22.

86. Mignone, F., et al., Untranslated regions of mRNAs. Genome Biol, 2002. 3(3): p. REVIEWS0004.

87. Rhind, N., et al., Comparative functional genomics of the fission yeasts. Science, 2011. 332(6032): p. 930–6.

88. Lynch, M., D.G. Scofield, and X. Hong, The evolution of transcription-initiation sites. Mol Biol Evol, 2005. 22(4): p. 1137–46.

89. David, L., et al., A high-resolution map of transcription in the yeast genome. Proceedings of the National Academy of Sciences, 2006. 103(14): p. 5320–5325.

90. Chen, C.-H., et al., The plausible reason why the length of 5’ untranslated region is unrelated to organismal complexity. BMC Research Notes, 2011. 4: p. 312.

