## Supplementary_Tables_Figs for "The Patterns of Alternative TSS Usage Explain the Highly Heterogeneous Landscape of 5’UTR Lengths in Eukaryotes"

### Table S1. Mapping statistics of CAGE datasets in yeast species

| sample | total | overall alignment rate | unique mapped | unique mapped rate |
|---|---|---|---|---|
| ScerAr.1 | 39938546 | 94.93% | 27771775 | 69.54% |
| ScerAr.2 | 44730229 | 94.75% | 31569089 | 70.58% |
| ScerDD.1 | 29567980 | 95.04% | 14434829 | 48.82% |
| ScerDD.2 | 24919356 | 95.42% | 11560704 | 46.39% |
| ScerDS.1 | 35161190 | 95.66% | 22129261 | 62.94% |
| ScerDS.2 | 36682729 | 95.60% | 22315465 | 60.83% |
| ScerGa.1 | 45714017 | 95.40% | 22366435 | 48.93% |
| ScerGa.2 | 39070001 | 94.80% | 20978144 | 53.69% |
| ScerGl.1 | 42056614 | 95.36% | 21851098 | 51.96% |
| ScerGl.2 | 38553431 | 95.18% | 20938226 | 54.31% |
| ScerH2.1 | 41669661 | 95.41% | 18871366 | 45.29% |
| ScerH2.2 | 17907214 | 95.22% | 8425339 | 47.05% |
| ScerHS.1 | 43747268 | 95.67% | 19614393 | 44.84% |
| ScerHS.2 | 25762666 | 95.73% | 9090311 | 35.28% |
| ScerNa.1 | 34638352 | 95.11% | 19977331 | 57.67% |
| ScerNa.2 | 19037627 | 94.72% | 10363229 | 54.44% |
| ScerYP.1 | 29740675 | 95.17% | 17346210 | 58.32% |
| ScerYP.2 | 30005902 | 95.88% | 20196591 | 67.31% |
| Spar.1 | 41471738 | 89.26% | 21260671 | 51.27% |
| Spar.2 | 43638383 | 83.48% | 20380879 | 46.70% |
| S. castellii-1 | 36763702 | 96.25% | 29365574 | 79.88% |
| S. castellii-2 | 37560080 | 96.06% | 29540910 | 78.65% |
| Sch. pombe -1 | 41068801 | 90.87% | 32673739 | 79.56% |
| Sch. pombe -2 | 35638178 | 90.11% | 28826177 | 80.89% |

**Table S2.A list of omics data used in this study**

| Dataset | Species | Strain/Cell line | Condition | Growth phase | Temperature | CHX treatment | rRNA deletion | Type |
|---|---|---|---|---|---|---|---|---|
| PRJNA483730 | S. cerevisiae | BY4741 | Ar: Arrest, α factor (2.5 mM) for 45 min; add another 50 μL to 25 mL yeast for another 30 min | log-phase | 30°C | N.A | N.A | CAGE |
| PRJNA483730 | S. cerevisiae | BY4741 | YP: YPD | log-phase | 30°C | N.A | N.A | CAGE |
| PRJNA483730 | S. cerevisiae | BY4741 | DD: 1 mM MMS for 1 h | log-phase | 30°C | N.A | N.A | CAGE |
| PRJNA483730 | S. cerevisiae | BY4741 | DS: DSA, YPD medium for 48 h | log-phase | 30°C | N.A | N.A | CAGE |
| PRJNA483730 | S. cerevisiae | BY4741 | Ga: YP medium with 2% galactose | log-phase | 30°C | N.A | N.A | CAGE |
| PRJNA483730 | S. cerevisiae | BY4741 | Gl: YP medium with 16% glucose | log-phase | 30°C | N.A | N.A | CAGE |
| PRJNA483730 | S. cerevisiae | BY4741 | H2: Add $H_2O_2$ to early-log phase cells for a final concentration of 0.20 mM, 30 min | log-phase | 30°C | N.A | N.A | CAGE |
| PRJNA483730 | S. cerevisiae | BY4741 | HS: Heat shock from 30°C to 37°C, 1 h | log-phase | 30°C | N.A | N.A | CAGE |
| PRJNA483730 | S. cerevisiae | BY4741 | Na: Add NaCl for a final concentration of 1 M for 45 min | log-phase | 30°C | N.A | N.A | CAGE |
| PRJNA510689 | S. paradoxus | N17 | YPD | log-phase | 30°C | N.A | N.A | CAGE |
| PRJNA510689 | Sch. pombe | JB1171 | YPD | log-phase | 30°C | N.A | N.A | CAGE |
| FONTOM5 | Homo sapiens | K562 | RNA directly extracted from stock | N.A | 37°C | N.A | N.A | CAGE |
| FONTOM5 | Homo sapiens | HepG2 | RNA directly extracted from stock | N.A | 37°C | N.A | N.A | CAGE |
| PRJNA313018 | Mus musculus | NIH3T3 | DMEM | N.A | 37°C | N.A | N.A | CAGE |
| PRJNA313018 | Mus musculus | NIH3T3 | DMEM | N.A | 37°C | Yes | no | Polysome profiling |
| SRP187756 | S. cerevisiae | S288C | H2: Add $H_2O_2$ to early-log phase cells for a final concentration of 1.5 mM, 30 min | log-phase | 30°C | 100 μg/ml | N.A | Polysome profiling |
| SRP187756 | S. cerevisiae | S288C | YP: YPD | log-phase | 30°C | 100 μg/ml | N.A | Polysome profiling |

**Table S3. Number of ATU genes by setting the dominant TSS distance to 50 nt, 25nt or 0 nt**

| Condition | Total gene | >50 nt | >25 nt | >0 nt |
|---|---|---|---|---|
| Cell Arrest | 4981 | 142 | 228 | 2536 |
| DNA Damage | 4828 | 108 | 220 | 2566 |
| Diauxic Shift | 4835 | 191 | 359 | 3340 |
| Galactose | 5111 | 145 | 262 | 3898 |
| Gluctose | 4841 | 130 | 237 | 2484 |
| H2O2 | 4972 | 109 | 208 | 2421 |
| Heat Shock | 4803 | 188 | 340 | 2930 |
| NaCl | 5049 | 144 | 280 | 3523 |

**Table S4. Statistics of ATU and DE genes in different conditions comparing to YPD control**

| Condition | # Gene with ATU | #gene without ATU | # Gene with ATU and DE | # Non ATU Gene with DE | #total |
|---|---|---|---|---|---|
| Cell Arrest | 142 | 4839 | 77 | 628 | 4981 |
| DNA Damage | 108 | 4720 | 56 | 1254 | 4828 |
| Diauxic Shift | 191 | 4644 | 145 | 2269 | 4835 |
| Galactose | 145 | 4966 | 93 | 1935 | 5111 |
| Glucose | 130 | 4711 | 75 | 1090 | 4841 |
| $H_2O_2$ | 109 | 4863 | 57 | 746 | 4972 |
| Heat Shock | 188 | 4615 | 141 | 1774 | 4803 |
| Nacl | 144 | 4905 | 70 | 1469 | 5049 |

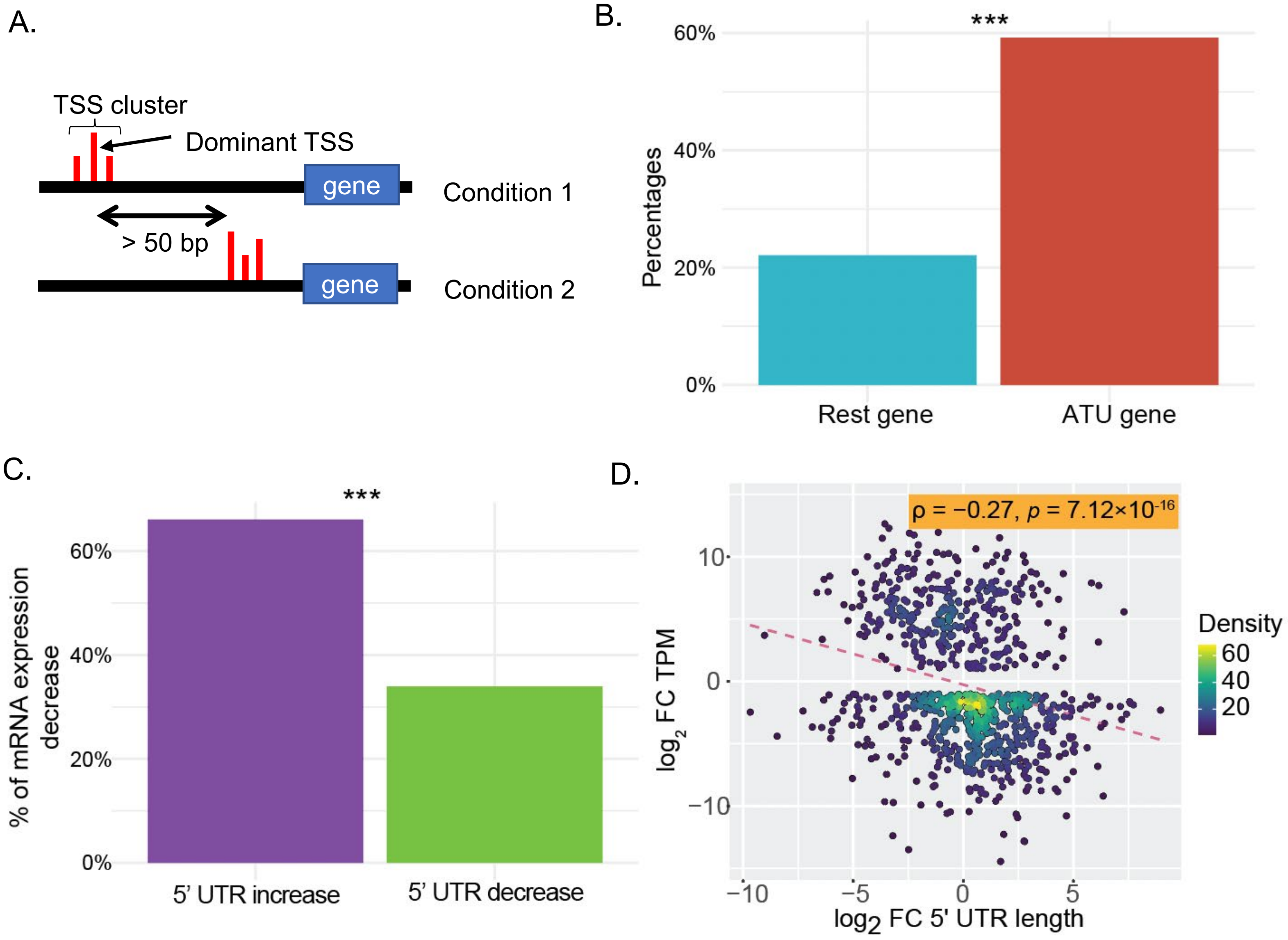


**Supplementary Figure S1. The percentages of significantly differentially expressed (DE) genes are significantly higher in ATU genes than in rest genes. (A)** An illustrative depiction outlining the definition of ATU: Firstly, the distribution of the two TSS clusters is statistically different as determined through the utilization of a paired Wilcoxon Test ($p < 0.05$); Secondly, the dominant TSS in each TSS cluster displays a certain distance (≥ 50 bp). **(B)** The percentages of DEG (FC ≥ 2) in ATU genes or all genes by comparing two human cell lines (HepG2 and K562). The X-axis shows different comparisons. The Y-axis shows percentages. ***: $p < 0.001$, $p$ value is from $\chi 2$ test. **(C)** % of transcripts expression decrease when 5' UTR length increases or decreases. ***: $p < 0.001$. The $p$ value is from a binomial test of the null hypothesis of equal numbers of genes in the two groups. **(D)** FC of 5' UTR length anticorrelates with FC of gene expression by comparing TSS maps of two human cells lines (HepG2 and K562); The X-axis shows FC of 5' UTR length ($log_2$), the Y-axis show FC of gene expression ($log_2$). The genes with significant changes of transcription level (FC ≥2) and ATU ($p$ <0.05, wilcox.test) were identified between two cell lines.

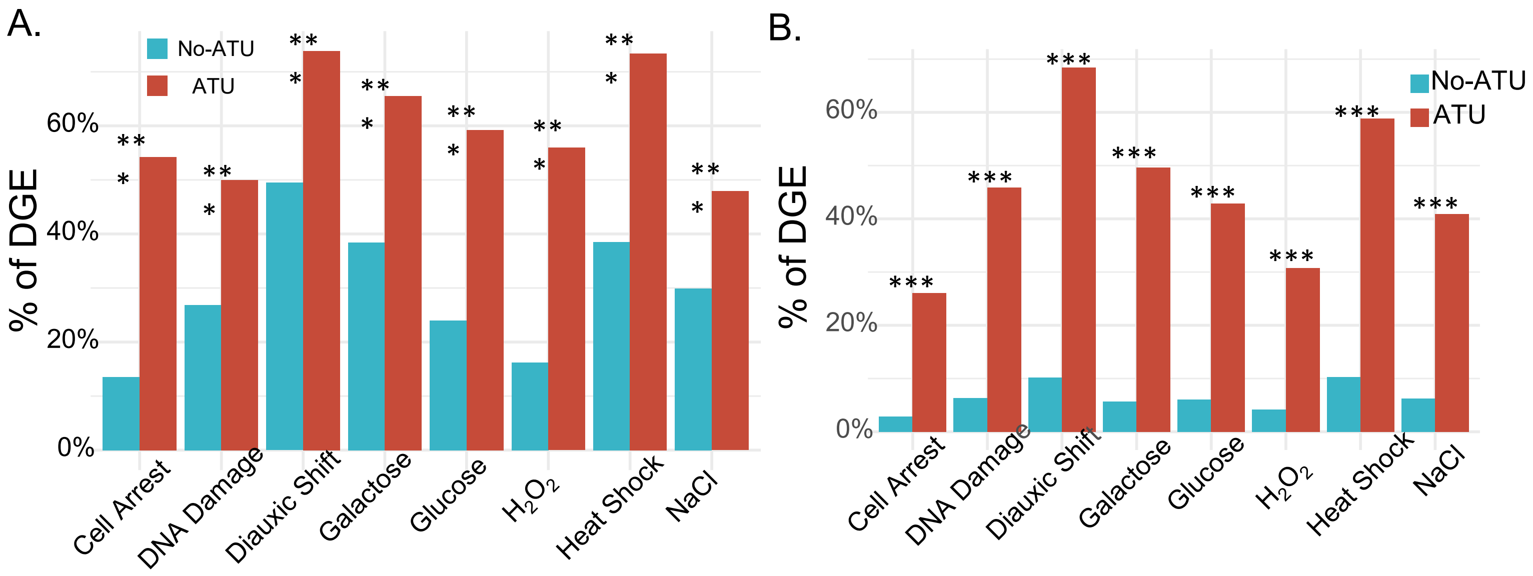


**Supplementary Figure S2. The percentages of significantly differentially expressed (DE) genes are significantly higher in ATU genes than in the rest genes. (A)** The percentages of significantly differentially expressed genes (DEG) (FC ≥ 2) in ATU genes ($p < 0.05$, the distance of dominant TSS ≥ 50nt, Materials and Methods) or non-ATU genes  the X axis is different treatment vs control condition (YP), the Y axis is the percentages of DGE genes. YP: YPD rich media, Cell Arrest: cell arrest by the alpha factor, DNA Damage: DNA damage. Diauxic Shift: diauxic shift. Galactose: carbon source, galactose, Glucose: fermentation by 16% glucose, $H_2O_2$: oxidative stress by $H_20_2$, Heat Shock: heat shock, NaCl: osmotic stress (NaCl). **(B)** Similar to A, except for the criteria of ATU genes which was set to $p < 0.05$, the distance of dominant TSS ≥ 0 nt. ***: $p < 0.001$.

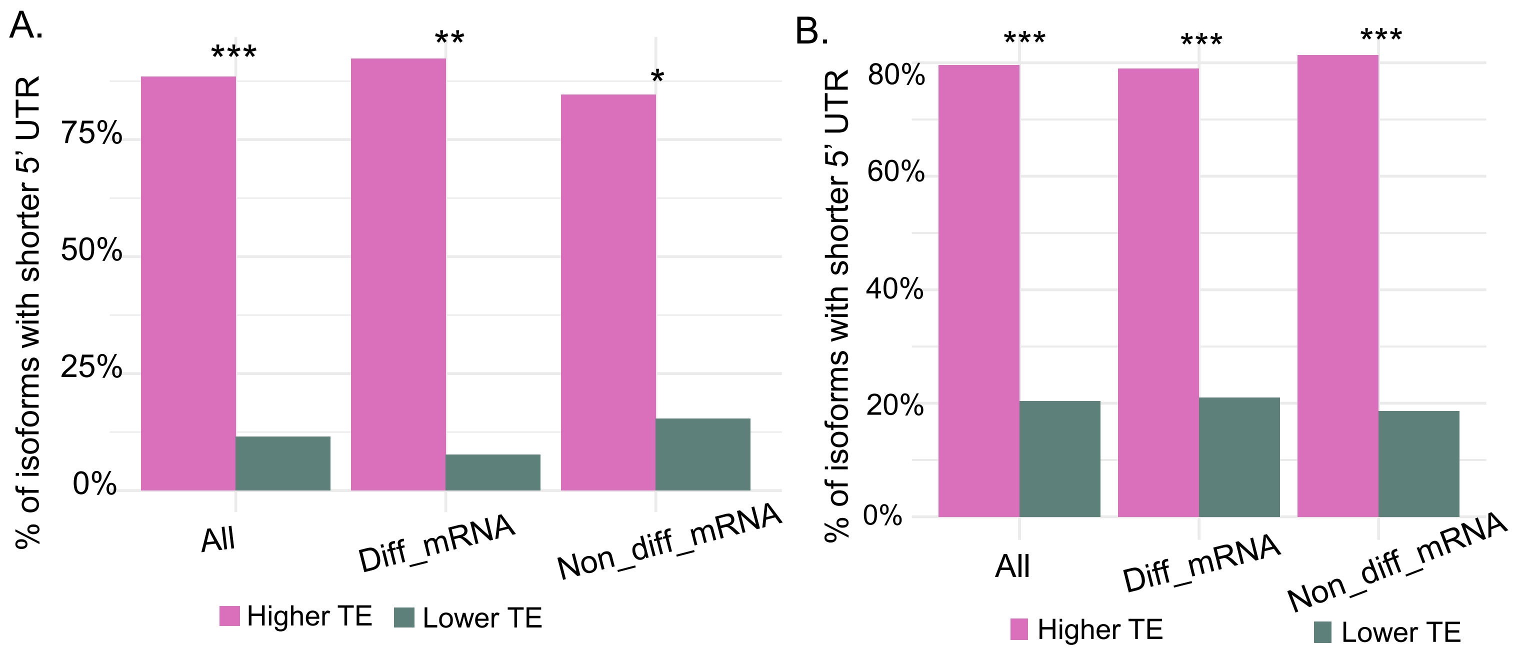


**Supplementary Figure S3. Gene with long 5'UTR tend to have a low TE and the effect is independent of mRNA level (A)**, Diff_mRNA" refers to the group of genes that there is significant difference in the abundance between mRNA isoforms (|mRNA2/mRNA1| ≥ 2). "Non_diff_mRNA" refers to the groups of genes which there is no significant difference in the abundance between mRNA isoforms (|mRNA2/mRNA1| < 2). Only genes with significant TE are compared (|TE2/ TE 1| ≥ 1.5), the data was from *S. cerevisiae*. **(B)** similar to (A) except for the data was from mouse fibroblasts (NIH3T3). *: $p < 0.05$, **: $p < 0.01$, ***: $p < 0.001$, binomial test.

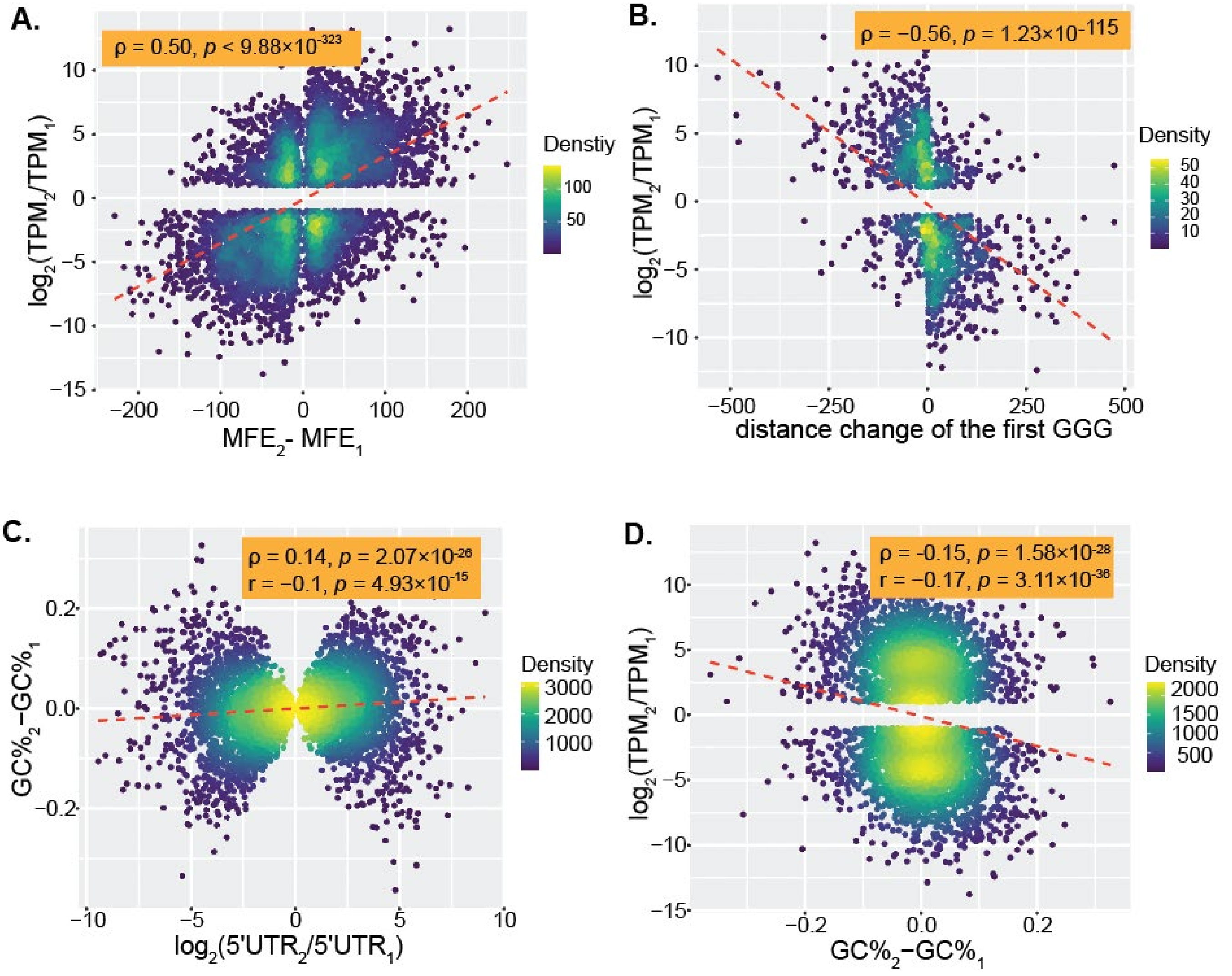


**Supplementary Figure S4. Correlation between different motif characteristic and peak TPM or changes of peak TPM ($log_2$). (A)** A positive correlation between changes of MFE [x = $MFE_2$ - $MFE_1$] and fold changes of transcript abundance (y = $log_2(TPM_2/TPM_1)$).
**(B)** A negative correlation between change of the distance of the first GGG to TSS (dGGG2-dGGG1) and fold changes of transcript abundance (y = $log_2(TPM_2/TPM_1)$ from transcription isoforms generated by the same gene. Each dot represents a pair of 5'UTR isoforms in S. cerevisiae.**(C)** Scatter plot showing a positive correlation between $log_2$ ratios of 5'UTR [*x* = $log_2(5'UTR_2/5'UTR_1)$] and changes of GC% content [y=$GC\%_2$-$GC\%_1$] from transcription isoforms generated by the same gene. **(D)** Scatter plot showing a negative correlation between changes of GC% content [x=$GC\%_2$-$GC\%_1$] and peak TPM values [*y* = $log_2(TPM_2/TPM_1)$] from transcription isoforms generated by the same gene.

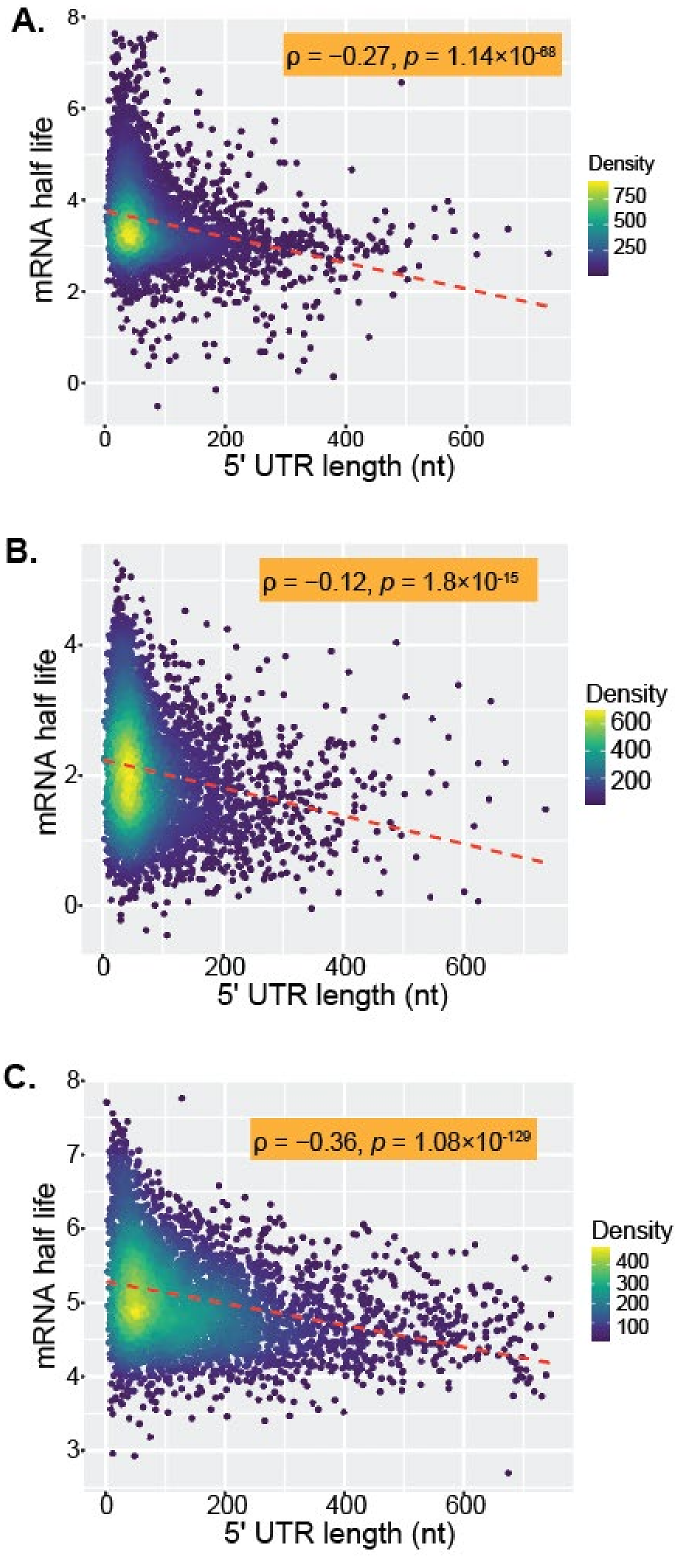


**Supplementary Figure S5. 5’UTR lengths is anticorrelated with mRNA half life.**
**(A)** Negative correlations between 5’UTR lengths and mRNA half life. X axis is the weighted 5’ UTR length and Y axis is the mRNA half life data from Gresham et al 2015. **(B)** Negative correlations between 5’UTR lengths and mRNA half life in *S. cerevisiae*. X axis is the weighted 5’ UTR length and Y axis is the mRNA half life data. The mRNA half data is from Chan et al 2018. **(C)** Negative correlations between 5’UTR lengths and mRNA half life in *Sch. pombe*. The mRNA half data is from Mata et al 2015.

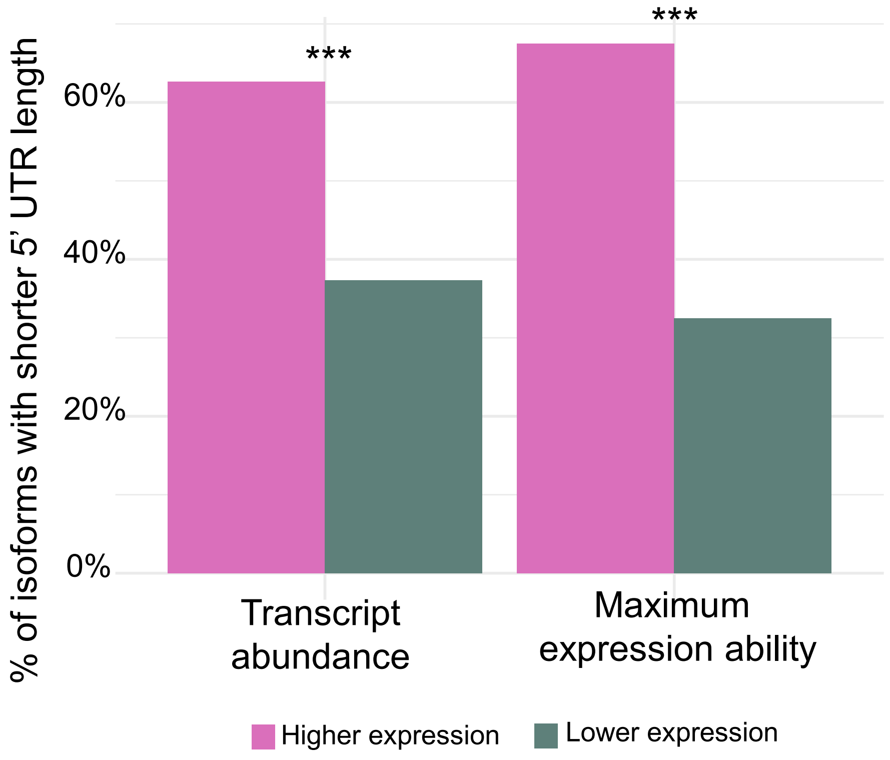


**Supplementary Figure S6. Transcript with short 5' UTR length tend to have high expression levels** Maximum expression ability: Peak expression of transcript isoform in nine conditions. ***: $p$ < 0.001, binomial test. Only transcript isoform pairs with significant different expression are compared ($|mRNA_2/ mRNA_1| \geq 2$).

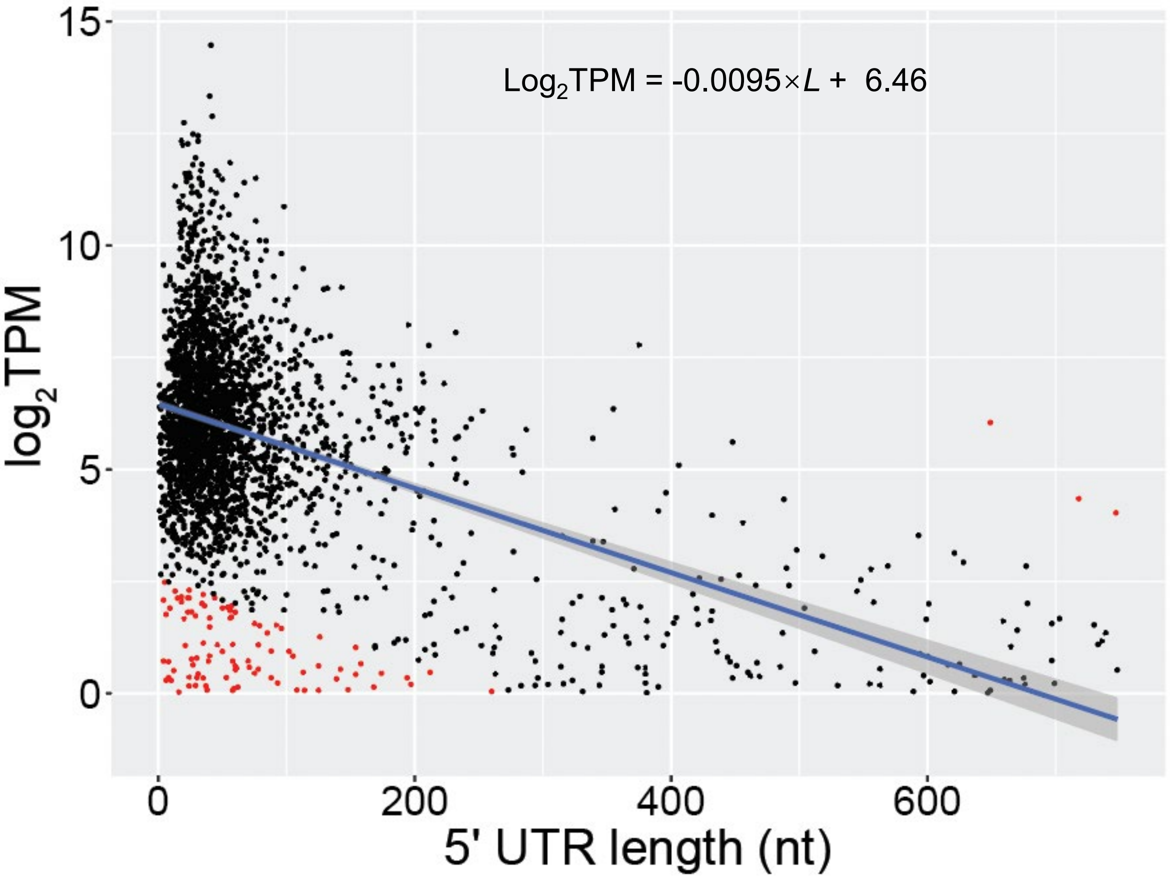


**Supplementary Figure S7. A simple linear model shows the relationship between 5' UTR lengths and peak expression level from gene with a single core promoter.** Each dot represent a gene and dots in red color were identified as genes with a non-optimal 5'UTR length or outlier genes.

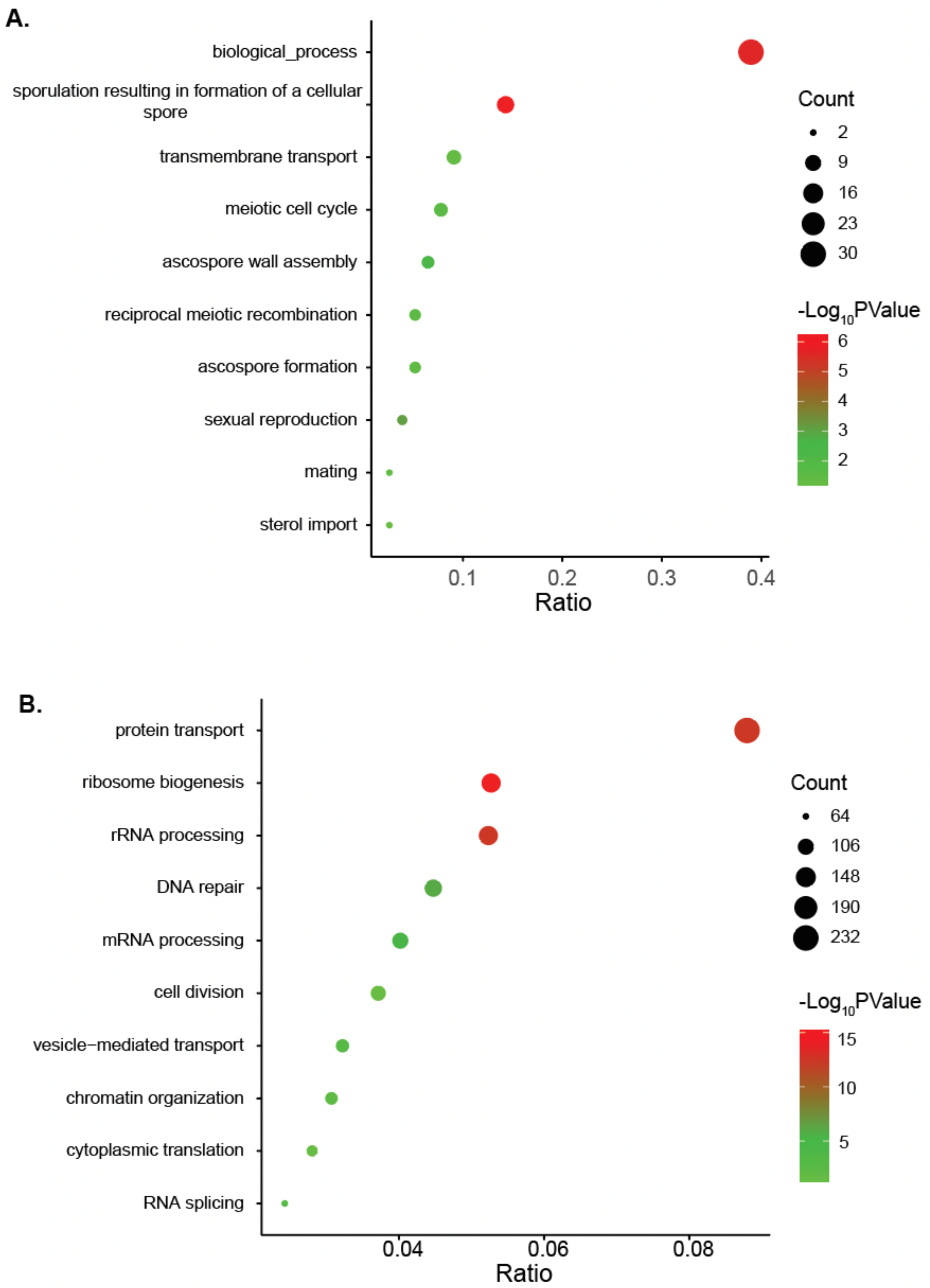


**Supplementary Figure S8. Functional enrichment of 5'UTR optimal and non-optimal genes. (A)** KEGG pathway enrichment of 5'UTR non-optimal genes **(B)** KEGG pathway enrichment of 5'UTR optimal genes**.**

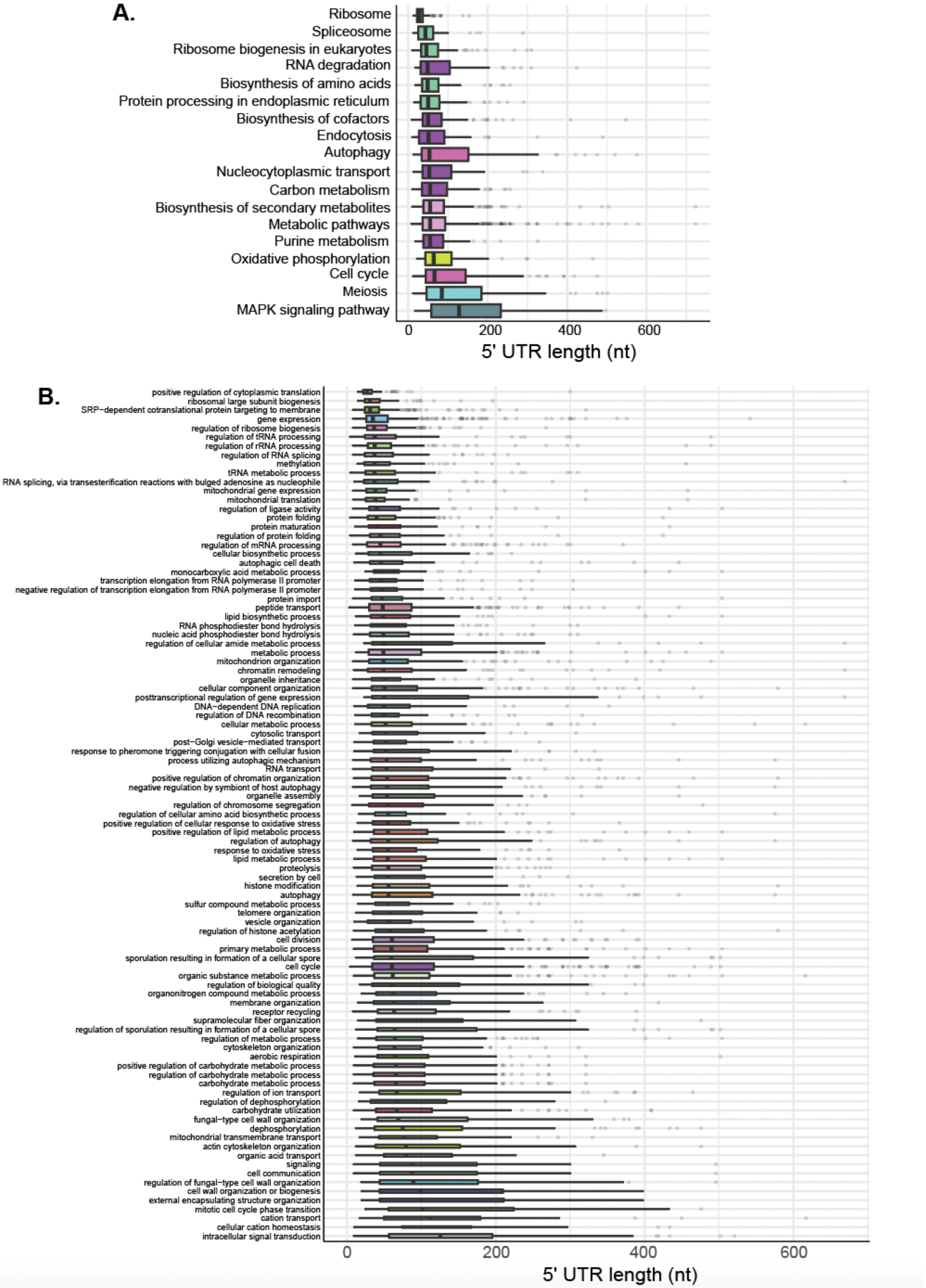


**Supplementary Figure S9. Distribution of 5'UTR length among different types of genes. (A)** Box plots showing large variations in 5′ UTR lengths among genes from different KEGG pathways. **(B)** Box plots showing large variations in 5′ UTR lengths among genes from different GO terms.

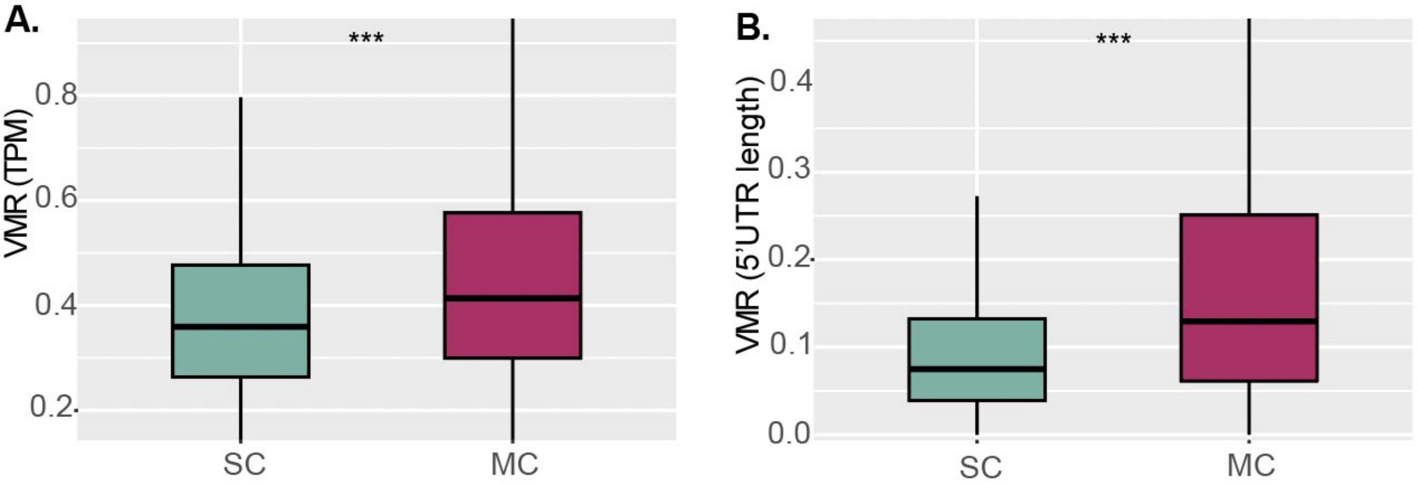


**Supplementary Figure S10. Higher variance of gene expression and 5'UTR length in MC genes than SG genes**. **(A)** Boxplot showing a much higher relative expression variance in MC genes than SC genes in *S. cerevisiae*. The relative variance, or variance-to-mean ratio (VMR) was calculated as the standard deviation of TPM values across nine growth conditions normalized by its mean. **(B)** Boxplot showing a much higher relative variance of 5' UTR length in MC (multiple-cluster) genes than SC (single-cluster) genes. The relative variance was calculated as the standard deviation of 5'UTR length values across nine growth conditions normalized by its mean. ***: $p$ < 0.001, Student's t-test.
